# A shared gene but distinct dynamics regulate mimicry polymorphisms in closely related species

**DOI:** 10.1101/2025.03.03.641230

**Authors:** Sofia I. Sheikh, Meredith M. Doellman, Nicholas W. VanKuren, Phoebe Hall, Marcus R. Kronforst

## Abstract

Sex-limited polymorphisms, such as mating strategies in male birds and mimicry in female butterflies, are widespread across the tree of life and are frequently adaptive. Considerable work has been done exploring the ecological pressures and evolutionary forces that generate and maintain genetic variation resulting in alternative sex-limited morphs, yet little is known about their molecular and developmental genetic basis. A powerful system to investigate this is *Papilio* butterflies: within the subgenus *Menelaides,* multiple closely related species have female-limited mimicry polymorphism, with females developing either derived mimetic or ancestral non-mimetic wing color patterns. While mimetic color patterns are different between species, each polymorphism is controlled by allelic variation of *doublesex* (*dsx*). Across several species, we found that the mimetic and non-mimetic females develop male-like color patterns when we knockdown *dsx* expression, establishing that *dsx* controls both sexual dimorphism and polymorphism. We also found that mimetic *dsx* alleles have unique spatiotemporal expression patterns between two species, *Papilio lowii* and *Papilio alphenor.* To uncover the downstream genes involved in the color pattern switch between both species, we used RNA-seq in *P. lowii* and compared the results to previous work in *P. alphenor*. While some canonical wing patterning genes are differentially expressed in females of both species, the temporal patterns of differential expression are notably different. Our results indicate that, despite the putative ancestral co-option and shared use of *dsx* among closely related species, the mimicry switch functions through distinct underlying mechanisms.

**AUTHOR SUMMARY:** Understanding how a largely shared genome can encode the potential to develop multiple morphs while simultaneously restricting this potential to one sex has long been of interest to evolutionary and developmental biologists. This phenomenon, called sex-limited polymorphism, is widespread, occurring in organisms like crustaceans, insects, fish, birds, and mammals. Recent empirical work has begun to identify the genes controlling the switch between phenotypes, but the differences between developmental programs leading to those phenotypes remain unclear. Here we use a classic example of sex-limited polymorphism – female mimicry in swallowtail butterflies – to compare how closely related species have evolved to use the same gene, *doublesex*, in the development of multiple female morphs. Using a combination of functional experiments, we show that despite the shared use of *doublesex*, the developmental genetics underlying sex-limited polymorphism have evolved to function quite differently between two species that last shared a common ancestor approximately 15 million years ago.

## INTRODUCTION

Despite the fact that sexes share the vast majority of their genomes, males and females can develop not only distinct phenotypes, but multiple alternate phenotypes suited to particular life histories and selective pressures (1–3). Sex-limited polymorphism – the ability of one sex to develop multiple discrete morphs – thus represents a striking case of adaptive phenotypic variation. This type of adaptation appears in disparate taxa and phenotypes (2), including the alternative mating strategies of male beetles and fish, where some males adopt sneaker or fighter morphs (4,5), and female color polymorphism in damselflies (6).

Studies characterizing the genetic control of sex-limited polymorphism have revealed that sex-specific trait variation can be achieved by encoding the polymorphism in the sex-linked genome. For example, in brood parasitic cuckoos, females exhibit distinct plumage morphs based on their W chromosome haplotype, while the homogametic males develop a single monochromatic pattern (7). In contrast, mapping studies have also identified many examples in which the genetic basis of sex-limited polymorphism is both autosomal and Mendelian, with allelic variation at a single locus acting as a switch between alternate phenotypes (for example, pigmentation in butterflies (8) and *Drosophila* (9)). In some cases, such switch loci function as supergenes, wherein multiple linked genetic elements contribute to the coordinated regulation of a complex phenotype in a simple Mendelian manner (2,10). This flurry of discoveries suggests that allelic variation at switch loci is sufficient to shape distinct developmental programs that produce multiple complex phenotypes.

The molecular mechanisms through which alternate alleles actually switch developmental programs to produce sex-limited polymorphism have been explored in only a handful of examples. In the brown anole, Feiner et al. (2022) suggest that alternate female color pattern morphs develop due to changes in cell migration behavior that may arise from alternate protein-coding sequences of *CCDC170* linked with estrogen receptor *ESR1* (11). In swordtail fish, the polymorphic false gravid spot in males, which mimics the pregnancy spot of female swordfish, is associated with alternate alleles of *kitlga* that have unique tissue and allele-specific expression patterns (12). Finally, in ruffs, three male mating morphs are distinguished by hormonal differences associated with distinct alleles at *HSD17B2* (13). Collectively, these examples illustrate that sex-limited polymorphism can develop by several potential mechanisms, including modifying gene expression patterns, altering protein function, or regulating hormone levels.

Beyond the limited number of functional case studies, explicit cross-species comparisons of the molecular and developmental control of sex-limited phenotype switching are missing. Yet, these comparisons are necessary to identify general principles on how developmental programs and gene regulatory networks (GRNs) evolve to produce alternate phenotypes. *Papilio* swallowtail butterflies offer a powerful natural system to investigate the evolutionary developmental genetics of sex-limited polymorphism in a comparative context. Several *Papilio* species have evolved female-limited mimicry polymorphism (FLMP), wherein males develop a single non-mimetic wing color pattern while females develop one of multiple discrete patterns, many of which mimic patterns of distantly related toxic species. In each case the switch between female patterns is controlled by allelic variation at an autosomal locus (Fig 1A). This female-limited polymorphism in *Papilio* butterflies is a classic example of a supergene (14–19).

In some species, such as *Papilio dardanus*, *Papilio polytes*, and *Papilio* alphenor, females can develop a male-like non-mimetic color pattern or one of several discrete mimetic patterns. In others, such as *Papilio memnon* and its relatives, none of the female morphs resemble the male color pattern. In *P. dardanus*, an inversion in the regulatory region of the transcription factor *engrailed* is associated with a diversity of mimetic female morphs (20,21). On the other hand, in the *Papilio* subgroup *Menelaides*, FLMP evolved through co-option of the conserved transcription factor *doublesex* (*dsx*) (14,18,19,22), which regulates sexual differentiation across insects (23,24). In addition to performing its ancestral function, alternate *dsx* alleles determine the adult female color pattern in at least six species (Fig 1A).

The evolutionary origins of mimetic *dsx* alleles across the *Menelaides* subclade remain unresolved, with current studies offering equivocal evidence for a single ancestral origin or multiple independent origins. Across the entire *Menelaides* subclade, species with FLMP have extremely divergent mimetic and non-mimetic *dsx* haplotypes. Mimetic *dsx* alleles across species have unique amino acid substitutions, distinct signatures of linkage disequilibrium, and varying structural features at the *dsx* locus (19,22,25). Within the *polytes* group, Zhang et al (2017) showed that species share a mimetic *dsx* haplotype, in addition to lineage-specific *dsx* haplotypes that produce additional mimetic female morphs (26). The mimetic (dominant) alleles share an inversion containing the *dsx* region and have both coding and non-coding differences from the non-mimetic alleles. In the *memnon* clade (*Papilio rumanzovia, memnon, and lowii*), no mimetic *dsx* alleles contain an inversion, and there are few to no coding sequence substitutions between the mimetic and non-mimetic haplotypes within a species (14,27). These divergent patterns of sequence evolution suggest independent co-option of *dsx* in different lineages (25). On the other hand, Palmer and Kronforst (2020) suggested that the differences across mimetic *dsx* alleles may be the result of allelic turnover, wherein ancestral alleles are continuously replaced with derived variants, thus obscuring evidence of shared ancestry among extant sequences (19). Whether *dsx* co-option evolved independently or ancestrally, the presence of a common switch gene across multiple species offers a critical look into how the same developmental machinery underlying sex-limited polymorphism have evolved since they last shared a common ancestor.

The functional basis of *dsx*-mediated mimicry has been extensively characterized in *Papilio polytes* and *P. alphenor*, with recent work highlighting the evolution of several novel cis-regulatory elements that autoregulate mimetic *dsx* alleles leading to a spike of *dsx* expression in mimetic females during early pupal development (28,29). This increased expression causes a suite of transcriptomic differences between mimetic and non-mimetic females throughout pupal wing development, ultimately resulting in the development of discrete adult female patterns (28,30,31). In *Papilio memnon*, *dsx* has been shown to control both mimetic and non-mimetic female color patterns, whereas the mimetic *dsx* allele causes mimetic abdominal pigmentation in females (32). Additionally, mimetic *dsx* does not appear to show a dramatic spike of expression during early pupal development like it does in *P. alphenor* mimetic females (32).

The combination of functional studies in *P. polytes, P. alphenor,* and *P. memnon* show that *dsx* is required for the mimicry switch and offers a general understanding of *dsx* expression in the context of wing color patterning among these species. However, what remains unknown is whether the *dsx* switch functions through the same developmental programs to specify color patterns in a sex-limited fashion. Characterizing these developmental programs requires a comparative analysis of gene expression in color pattern development across multiple species. To do this, we first tested the role of *dsx* expression in the adult color patterns across multiple species using RNAi, and then characterized spatial and temporal patterns of *dsx* expression and its downstream effects in *Papilio lowii* using a combination of immunohistochemistry and RNA-seq. By directly comparing *dsx* expression and signatures of genome-wide differential expression between *P. lowii* and *P. alphenor* (which last shared a common ancestor approximately 15 million years ago), we identify critical shared and unique elements of the wing developmental programs underlying the *dsx* mimicry switch and provide insight into the molecular mechanisms by which sex-limited polymorphisms evolve.

**Figure 1:**
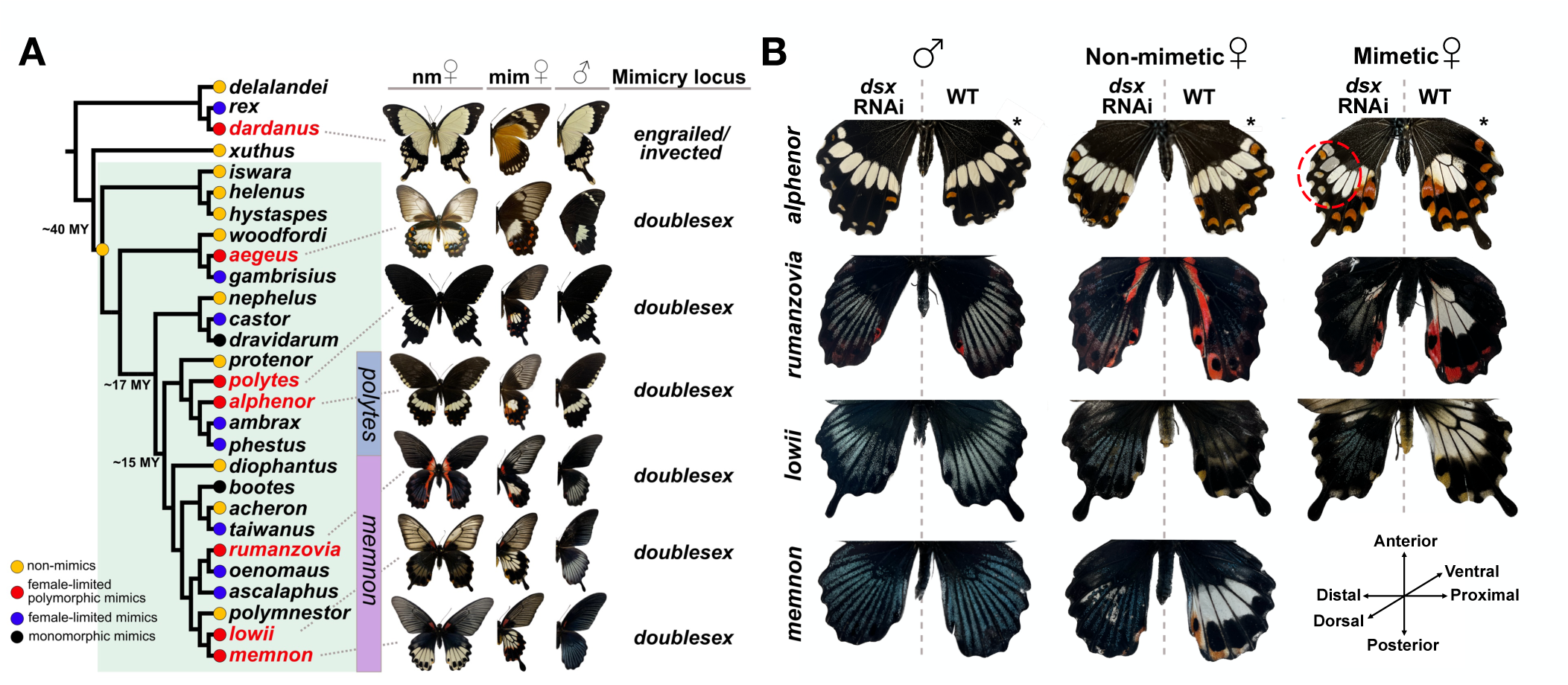
*Dsx*-controlled mimicry polymorphism in *Papilio* butterflies. **A)** Cladogram of a subset of *Papilio* species; *Menelaides* subclade highlighted in green (23 out of 56 known species and two species groups shown), with species relationships, divergence estimates, and mimicry state based on data from (19,33,34). “nm” refers to the non-mimetic female adult phenotype (homozygous for the recessive non-mimetic *dsx* allele), while “mim” refers to the mimetic phenotype. Not all alternate color patterns are shown for each species. **B)** *dsx* RNAi phenotypes on the dorsal surface of genotypically mimetic and non-mimetic females and male hindwing. Untreated wild-type (WT) wings serve as a control to compare with the knockdown phenotype within an individual. *Papilio alphenor* RNAi individuals are from (28). * Denotes individuals injected on the ventral surface and are displayed here as such. Orientation key applies to all images without an asterisk and is shown here for the treated wing relative to the midline. Additional experimental and control RNAi individuals in Supplementary Figure S1.

## RESULTS

### *dsx* has a conserved ancestral role specifying sexually dimorphic wing patterns

Genomic evidence has suggested that *dsx* controls the color pattern switch in all polymorphic *Menelaides* (14,18,19,22). However, the role of *dsx* in wing color patterning has only been examined in three species. In *P. alphenor* and *P. polytes,* knockdown of *dsx* expression in mimetic females results in a mosaic wing pattern resembling the non-mimetic female or male color pattern, whereas in *P. memnon*, mimetic females clearly converted to the male form in the absence of *dsx* (18,28). To confirm these results and test *dsx*’s role in wing color patterning in additional polymorphic species, we used an RNAi electroporation approach to knockdown *dsx* expression in the hindwing of three species in the *memnon* group: *P. rumanzovia, P. lowii*, and *P. memnon*. In contrast to the *polytes* group, polymorphic species in the *memnon* group display strong sexual dimorphism between both female morphs and males (Fig 1A). In these three species, RNAi caused females to develop a male-like color pattern (Fig 1B): treated regions of mimetic and non-mimetic females recovered a mosaic of the male-like adult color pattern in terms of both blue structural color and spatial organization of those scales across the wing. These results align with recent findings from *dsx* RNAi in *P. memnon* (32). *dsx* may be required for sexually dimorphic patterning in *P. alphenor* as well, however *dsx* RNAi in non-mimetic females has very minor and subtle effects on the adult color pattern, and non-mimetic females closely resemble males (28). Males of species in the *Memnon* group with mimetic or non-mimetic genotypes had slightly fewer blue scales in the *dsx* siRNAs injected region compared to both their WT wing and control males injected with PBS (Supplementary Figure S1), although the patterning and scale color were not particularly different. Altogether, these results suggest that *dsx* performs a conserved function in controlling sexual dimorphism, while the different *dsx* alleles control the development of distinct female color patterns.

### DSX expression prefigures mimetic color patterns but varies between species

After testing the role of *dsx* in female wing patterning, our second approach to assessing parallelism in the *dsx* mimicry switch was characterizing the expression patterns of DSX protein in the developing *P. lowii* hindwing. Previous work in *P. alphenor* showed that mimetic DSX has a dynamic expression pattern across the wing throughout early to mid-pupal development: in the early stages, it is highly and uniformly expressed across the wing, but eventually resolves to prefigure the central white patch in mimetic females (28). We sought to compare these spatial expression patterns with those from *P. lowii* using antibody staining at key developmental stages. The stains revealed that *dsx* is enriched in the hindwing of *P. lowii* mimetic females similarly to *P. alphenor*, however, its expression is spatially restricted from early-on during the fifth instar larval stage and remains localized throughout early to mid-pupal development (Fig 2A). This expression pattern prefigures the light bands that appear in the medial regions extending from the distal to proximal ends and toward the anterior end in mimetic females. Specifically, DSX is enriched in the scale precursor and socket cells from early pupal development. In mimetic females of *P. alphenor,* DSX is present in scale precursor, socket, and epidermal cells across the hindwing during early pupal development, and then gradually becomes spatially restricted to only the scale and socket cells that constitute the white patch in adult mimetic females (Fig 2A). Interestingly, DSX is also expressed in a spatially specific pattern in non-mimetic *P. lowii* females (Supplementary Figure S2), which is not the case in *P. alphenor* non-mimetic females (28).

In addition to spatial expression patterns, we quantified *dsx* expression in *P. lowii* using bulk RNA-seq of hindwings from late larval through late pupal development, spanning the window of color pattern specification (late larval to mid-pupal; L5-P6) and pigmentation (late pupal development; P9). We generated RNA-seq for *P. lowii* individuals homozygous for one *dsx* allele but included some heterozygous individuals for additional comparison (Supplementary Table S2 and Fig S4-S6). We compared these results to an existing and similar dataset in *P. alphenor* (NCBI SRA BioProject PRJNA882073 from ref. (28). Due to highly divergent *dsx* alleles and genome-wide heterozygosity, we assembled a long-read Nanopore genome sequence for *P. lowii* (Supplementary Table S3), and quantified gene expression for each species using its respective reference genome and annotation to avoid biasing quantification results. In *P. alphenor*, *dsx* is lowly expressed in all groups except mimetic females, where it undergoes a drastic and statistically significant increase of expression peaking at 2 days post-pupation. In contrast, we found that *dsx* is lowly expressed in all *P. lowii* groups (Fig 2B; Supplementary Fig S7). In mimetic females, *dsx* showed a pulse of expression during early pupal development, like in *P. alphenor*, however the difference in expression between mimetic and non-mimetic females was not statistically significant at any stage (overall FDR < 0.01).

**Figure 2:**
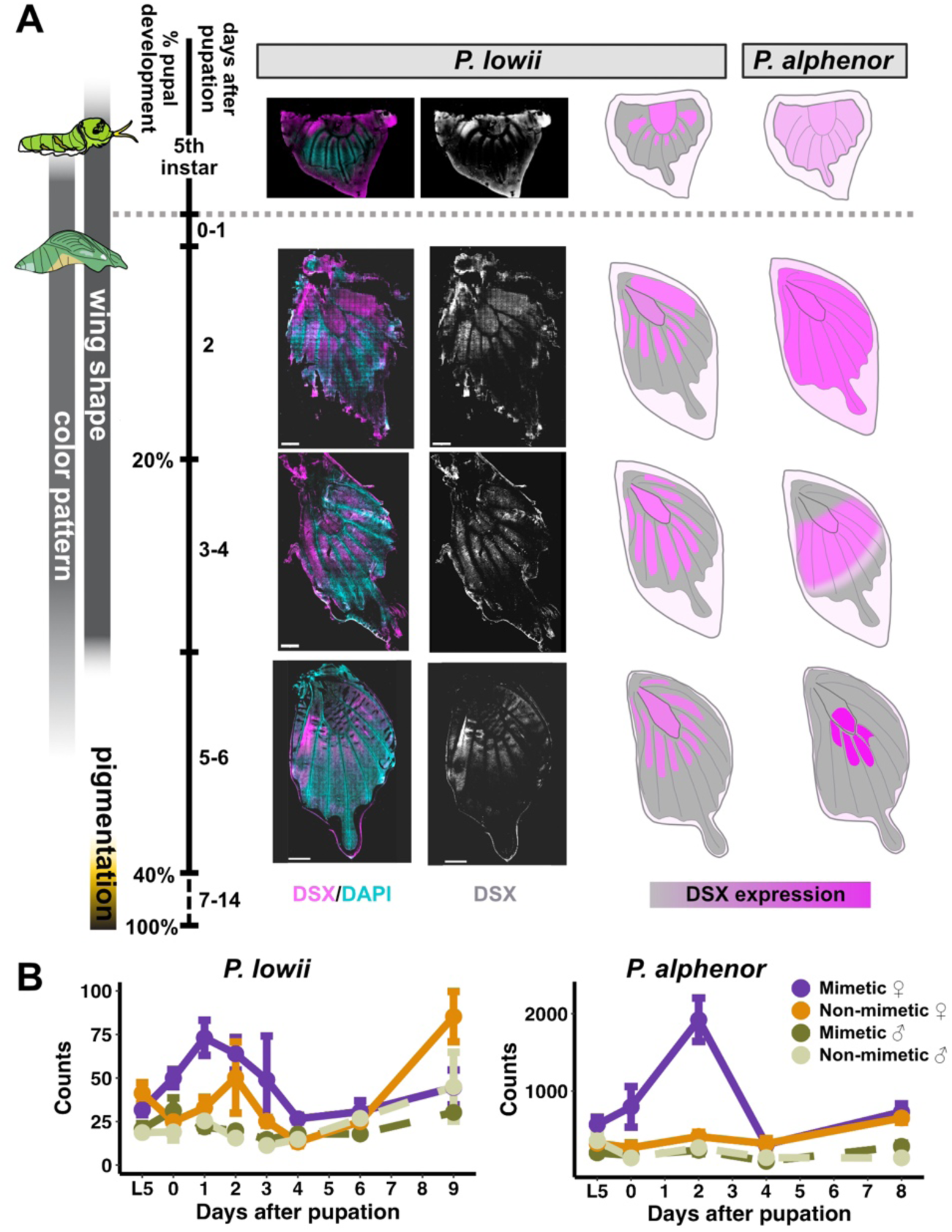
DOUBLESEX expression patterns in the hindwing of *P. lowii* and *P. alphenor*. **A**) Left: Anti-DSX staining from fifth-instar larval through mid-pupal development in hindwings from *P. lowii* mimetic female. Right: Cartoon schematic of DSX expression in *P. lowii* antibody stains and known patterns in *P. alphenor* at the same developmental time points (*P. alphenor* schematic modified from (28)). For additional staining images for *P. lowii* and late larval *P. alphenor*, see Supplementary Figures S2 and S3, and (28). Color pattern includes cell fate specification and differentiation, while pigmentation involves biosynthesis and transport of pigments. **B)** *dsx* normalized quantifications across development in four groups of *P. lowii* (left) and *P. alphenor* (right).

### Unique transcriptomic signatures of the *dsx* mimicry switch

Given the differences in mimetic DSX expression between *P. lowii* and *P. alphenor*, we sought to characterize how the different *dsx* alleles alter expression of downstream genes and gene regulatory networks to execute the mimicry switch in each species. First, we used two complementary approaches to identify differentially expressed genes (DEGs) in the RNA-seq datasets generated here for *P. lowii* and previously for *P. alphenor* (28). We identified DEGs between mimetic and non-mimetic females at each stage using DESeq2. We found 1,399 and 811 stage specific DEGs in *P. lowii* and *P. alphenor*, respectively. In *P. lowii*, divergent expression between mimetic and non-mimetic females was concentrated during late larval (L5) and late pupal development (9 days post pupation; P9). On the other hand, in *P. alphenor* almost all DEGs were found during early pupal development, coinciding with the spike of *dsx* expression at P2 (Fig 3A; right). Moreover, the majority of stage specific DEGs in *P. lowii* were upregulated in mimetic females, whereas in *P. alphenor* the opposite trend was observed. Additionally, the overall patterns of gene expression across development varied considerably (Fig 3A; left). While DESeq2 tested the effect of sex and *dsx* genotype on gene expression at each stage, we additionally tested for genes with distinct expression trends throughout the developmental time series using MaSigPro. In total, 571 and 679 genes exhibited unique longitudinal expression patterns (overall FDR < 0.01) in mimetic *P. lowii* and *P. alphenor* females, respectively (Fig 3B). All DEGs for each species are listed in Supplementary Tables S4 and S5.

**Figure 3:**
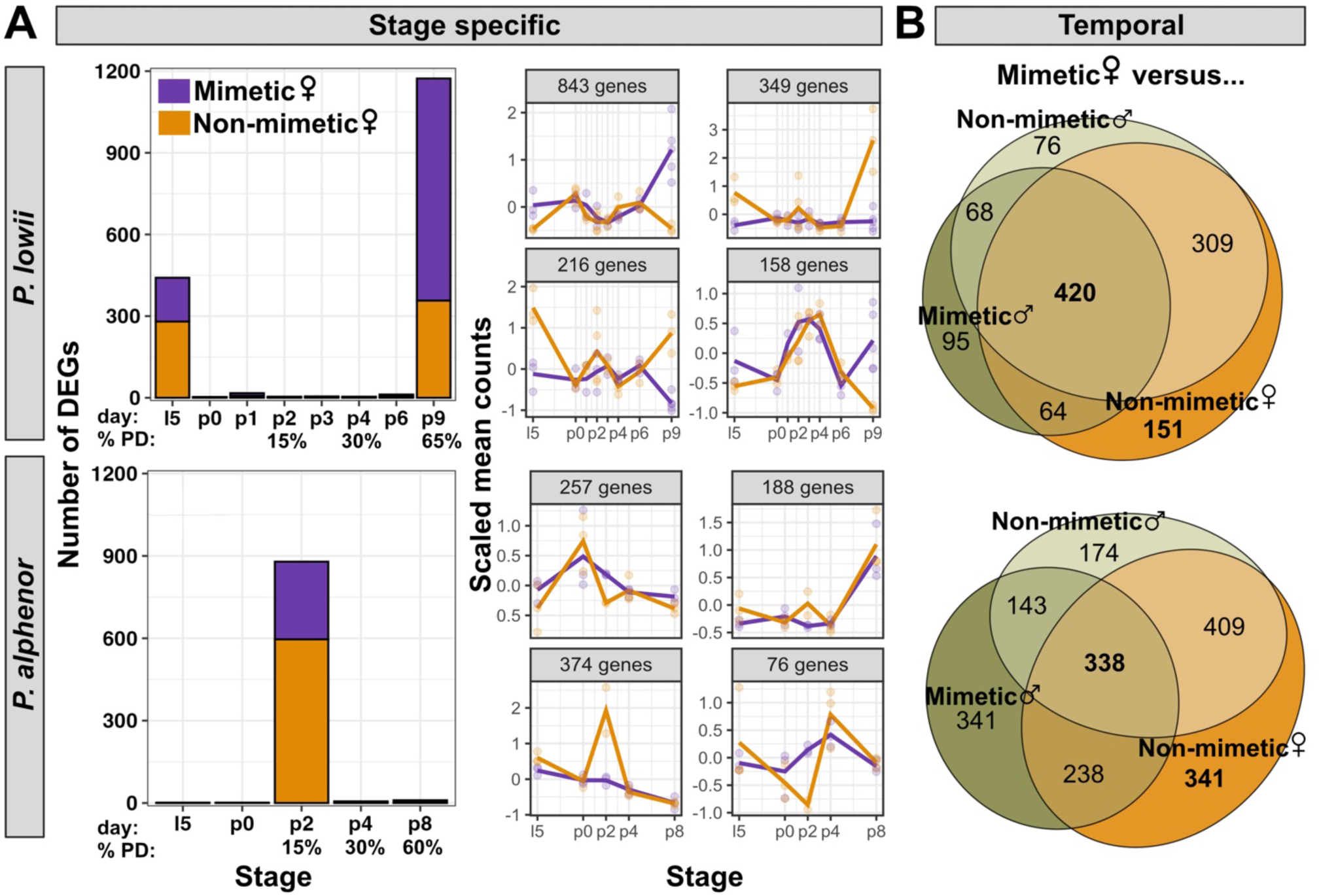
Differential gene expression underlying the mimicry switch in *P. lowii* and *P. alphenor*. **A)** Left: Genes differentially expressed and upregulated in either mimetic or non-mimetic females at each developmental stage. Day refers to the number of days after pupation and are directly comparable between both species because they share total pupal development (PD) time. Right: mean expression profiles of four largest clusters of genes in each species identified using DESeq2. **B)** Genes with significantly different temporal expression profiles in mimetic females relative to the other sex-genotype groups. The intersection represents genes unique to mimetic females, while the mimetic vs. non-mimetic female comparison represents genes that are unique to the non-mimetic females.

### Parallelism at the effector gene level

While the patterns of differential expression across development are variable and reflect potential differences in *dsx* regulation, the same downstream effector genes may be involved in both mimicry switches. Given the putatively ancestral co-option of *dsx* as a switch and close relatedness of the two species, we expected to find strong overlap between DEGs. In particular, due to the differences in the pigments and phenotypes of the adult female patterns, we expected to identify many shared pleiotropic transcription factors involved in cell fate specification instead of effector or terminal genes (for example, ones involved in pigment synthesis, transport, and deposition). To test this, we looked for overlap in the lists of DEGs using a one-to-one reciprocal best hit approach to identify orthologous genes between the two species; 90% (1772/1970) and 84% (1246/1490) of DEGs were assigned orthologous gene IDs in *P. lowii* and *P. alphenor*, respectively. Using these data, we identified genes that were significantly DE in both species, and ones that were uniquely DE within each species. We found that 18% of *P. lowii* DEGs were also differentially expressed in *P. alphenor* (Fig 4A; Supplementary Table S6). We performed a Fisher’s Exact Test based on genes that were differentially expressed in both species, unique to either species, or present in both transcriptomes but not differentially expressed, and found a statistically significant association between DEGs in both species (p = 3.6e-6).

We next wanted to investigate if known transcription factors (i.e. regulatory genes) or color patterning genes were unique or shared between the two mimicry switches. Among genes that were uniquely DE in each species, several are key transcription factors and morphogens of conserved signaling pathways that have been repeatedly co-opted in wing color patterning (35–38). For example, three of the *P. lowii* specific DEGs, *Notch, frizzled2* and *dishevelled* (a receptor for Notch signaling, and receptor and transducer for Wnt signaling, respectively), all have ancestral functions in cell-cell communication for cell fate specification and segment polarity (39,40). Secondarily, *Notch* has been co-opted into the wing developmental program to specify vein patterning and midline color patterns (37), while *frizzled2* receives *WntA*, which can act to prefigure and specify boundaries of adult color patterns (41). Interestingly, while these particular Wnt signaling components are not differentially expressed in *P. alphenor*, other key players are, including the regulator *shaggy*. Additionally, the *dsx* mimicry switch in *P. alphenor* appears to be mediated by several transcriptional co-regulators and targets like *yorkie* and *decapentaplegic* (28), which are not differentially expressed in *P. lowii*. Interestingly, we found that though *engrailed* was DE in only *P. alphenor*, preliminary immunohistochemistry data suggests that it functions similarly in *P. lowii* (Supplementary Figure S8).

Several of the shared genes also play key roles in early wing development. For example, the RNA-binding protein gene *couch potato* is enriched in sensory organ precursor pIIa daughter cells, which are destined to become scale and socket cells, as well as already dividing or divided scale cells in wings of the butterfly *Bicyclus anynana* (42). Similarly, c*rossveinless-c* encodes a BMP-binding protein that is required for wing vein morphogenesis (43), and *wings apart* inhibits the transcriptional repressor *capicua* in order to promote tissue growth and patterning, including of insect wings (44). Overall, among the 326 DEGs that were shared in the mimicry switch between both species, we identified functional groups using COG categories assigned to each gene by eggNOG (Fig 4B). In general, enrichment for translational processes and signal transduction mechanisms suggests that in both species, mimetic *dsx* likely has a drastic effect on cell specification and not the downstream/terminal processes in wing phenotype development.

### Parallelism at the GRN level

Finally, we aimed to compare and contrast the GRNs involved in the *dsx* mimicry switches using weighted gene co-expression network analysis (WGCNA). Each module was assigned a color at random and represents clusters of genes with strongly correlated expression profiles, thus potentially corresponding to biologically functional units (45). We found 23 and 31 modules of co-expressed genes in *P. lowii* and *P. alphenor*, respectively. After identifying modules, we tested for correlations between each module with various traits (stage, sex, genotype, and group), as well as modules that were significantly enriched with DEGs. In *P. lowii*, we found 13 modules that were significantly enriched or deficient in DEGs (Benjamini–Hochberg corrected FET p-value < 0.05). Of these, four modules were significantly correlated with mimetic females, that is, the module’s gene expression patterns were significantly associated with the group (Benjamini–Hochberg corrected FET p-value < 0.01; Supplementary Fig S9). In *P. alphenor*, 17 modules were significantly either enriched or deficient in DE genes, of which two were significantly negatively correlated with mimetic females (Supplementary Fig S10 and Supplementary Table S8). We first examined the most common processes from the top Gene Ontology (GO) terms of the first four modules that were most significantly enriched with DEs in each species. We found that modules most enriched for DEGs in each species were associated with some shared GO terms, including translation, protein localization, and metabolism (Fig 4C).

Next, we sought to compare whether the network modules were preserved across species and identify orthologous modules. Using the ModulePreservation function from WGNCA, we found that, generally, the overall network topologies (relationships, connectivity, and expression patterns of genes) were highly preserved between species. This is expected because the networks and modules for each species were generated from expression data from hindwing tissue, and the biological processes involved in wing development are likely to be conserved. However, despite this preservation in network topologies, the module compositions were considerably different in terms of gene membership between species, and we therefore were only able to identify orthologous modules for six out 24 *P. lowii* modules using a combination of the preservation score and gene membership count (Supplementary Fig S11). Of these orthologous modules, three were similarly associated with DEGs between species (one enriched and two deficient). The orthologous module enriched in both species for DEGs (green) had GO terms associated with toll signaling (Fig 4C). As part of its role in immune defense, toll signaling is involved in melanization, and this process is also used to control stripe patterning in *Bombyx mori* caterpillars (46). Additionally, several toll signaling genes are upregulated in other butterfly wing patterns, including eyespots in *B. anynana* (47), wing spots in *Pieres candida* (48), and red spots in mimetic *P. polytes* females (49).

**Figure 4:**
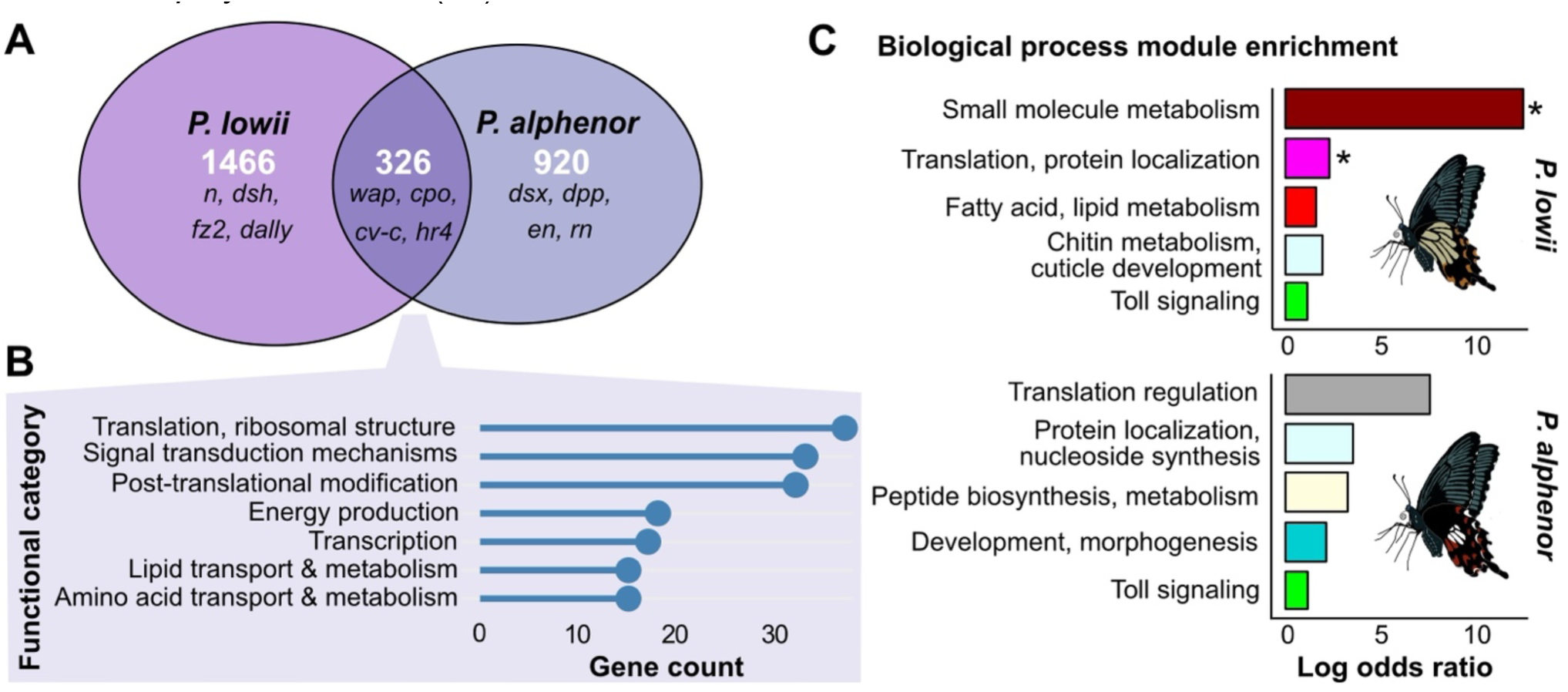
Developmental programs and pathways associated with the mimicry switch. **A)** Euler diagram of overlapping genes that are DE between mimetic and non-mimetic females of each species. Example of candidate genes based on literature listed for each set. Full results can be found in Supplementary Table S6 **B)** Top COG functional categories (> 15 genes) assigned to the overlapping DEGs in panel A. **C)** Gene ontology term enrichment for the top four co-expressed gene modules containing the highest number of DEGs in each species, and the single most strongly correlated module between both species. Colors reflect the module color assigned by WGCNA (Supplementary Fig S6-8). *Denotes modules that were also significantly correlated with the mimetic female group.

## DISCUSSION

Here we compared the role of *dsx* in orchestrating the development of discrete, complex female morphs among closely related *Papilio* swallowtail butterflies. This system presents a unique case of parallelisms at multiple scales – the phenotypic state of mimicry; sexual dimorphism and female polymorphism; the switch gene dictating the adult female color pattern; regulation of switch gene expression; and potentially its downstream consequences on developmental mechanisms for wing patterning. To begin dissecting the different levels of parallelism, we first characterized the role of *dsx* in wing color patterning of polymorphic species. Our results show that sexual dimorphism in wing patterning is governed by *dsx,* consistent with its role in controlling insect sexual dimorphism (23,24). Importantly, in addition to *dsx*’s role in sexually dimorphic wing patterning, it is allelic variation at *dsx* that determines which color patterns females develop by switching between alternate wing pattern developmental programs.

To further characterize how the *dsx* switch gene functions across species, we focused additional functional experiments on one species*, P. lowii*, due to its evolutionary distance from the well-studied species, *P. alphenor*. In comparing *dsx* expression from our experiments in *P. lowii* with previous work in *P. alphenor* (28), our results suggest that the mimetic alleles are expressed in notably different patterns, both in terms of their absolute levels and their spatial patterns across the developing wing. Moreover, the non-mimetic *dsx* alleles appear to have diverged in their expression between species. These results may suggest that *dsx* was convergently co-opted in mimicry polymorphism evolution two or more times. Alternatively, this likely represents cis-regulatory divergence in the *dsx* supergene following co-option in the common ancestor, in concordance with the idea of allelic turnover from an ancestral mimetic *dsx* in *Papilio* as proposed by Palmer and Kronforst (19). Future work uncovering the regulatory landscapes of different *dsx* alleles across *Menelaides* is necessary to determine the shared and lineage-specific repertoire of CREs that instruct differential spatiotemporal DSX expression both between morphs within a species, as well as among the mimetic female morphs of multiple polymorphic species.

Beyond the switch gene itself, we examined transcriptomic changes due to mimetic *dsx* alleles in *P. alphenor* and *P. lowii* in terms of the genes and GRNs involved in the development of mimetic female patterns. We observed surprising differences in the developmental windows at which gene expression was significantly altered between mimetic and non-mimetic females, suggesting that unique developmental trajectories underlie the mimetic female phenotype across these species despite the shared use of *dsx* and their close evolutionary relatedness. Nonetheless, we also identified some critical wing cell fate specification and patterning genes that were differentially expressed in both species, suggesting that both species may share some proximate functional mechanisms of mimetic wing patterning by *dsx*. These genes may be directly regulated by *dsx* or represent critical nodes of wing developmental networks that facilitate phenotype switching among females. Interestingly, roughly a quarter of the genes that were differentially expressed in both species are also differentially bound by DSX in *P. alphenor* mimetic females compared to non-mimetic females (29), and thus represent promising genes to target for future functional characterization.

Overall, there appear to be considerable lineage-specific differences despite both species being closely related and using the same switch gene for FLMP. While constraints exist in wing GRNs (50), the GRNs themselves are dynamic throughout development (51). The opposing windows of differential expression that we observed between *P. lowii* and *P. alphenor* suggest that DSX may be modifying expression of distinct elements of wing developmental GRNs. Additionally, wing shape, pattern and color differ between species and this could be responsible for some of the observed molecular differences. For example, in *P. memnon* and *P. alphenor*, tails are polymorphic and only develop in mimetic females, whereas the tail is a monomorphic structure in *P. lowii*. Similarly, abdominal pigmentation is only observed in mimetic females of *P. memnon*. These differences suggest the presence of species-specific modifier genes. While supergene theory emphasizes the physical linkage of functionally related genetic elements (10,52), unlinked modifiers—genes not physically linked to the switch locus—can epistatically interact to enhance mimicry (53). The unique DEGs we identified between species likely represent some of the species and morph-specific modifier genes required to execute each mimicry switch.

Another potential explanation for the observed divergence between *P. lowii* and *P. alphenor* may be developmental systems drift, wherein the genetic and regulatory basis of homologous traits can diverge even among closely related species without changing the trait itself (54). One of the most obvious cases of phenotypic invariance coupled with highly divergent developmental underpinnings is sex determination and differentiation across a multitude of taxa (55). For example, in dipteran flies, the sex determination hierarchy diverges in upstream signals (*Sxl* in *Drosophila* vs. *F* in *Musca*) while the downstream effector (*dsx*) is conserved, maintaining the same developmental outcome despite lineage-specific genetic interactions (54,56). In *dsx*-mediated mimicry, it is possible that an ancestral co-option of *dsx* was followed by lineage-specific drift in the developmental genetics underlying female polymorphism. This may be reflected by the considerable overlap yet differences in DEGs, and the observation that different genes in known wing specification and patterning pathways are DE within each species. Comparing the direct targets of mimetic DSX and genetic manipulations of DSX targets across species will help test the relative roles of species-specific modifiers and developmental systems drift in the developmental differences among sex-limited polymorphic species.

Finally, we note that bulk RNA-seq approaches to characterizing homology in molecular mechanisms and GRNs have several limitations. The loss of cellular-level resolution can obscure critical signals and differences between cell types (for example, transcriptomes of *dsx*-expressing cells during early and mid-pupal development vs. that of cells specified to become melanic) (57). On the other hand, the lack of conservation in gene expression patterns across species may not indicate functional divergence (57). Such cross-species comparisons may also suffer from technical challenges, including identifying orthologous genes and reference genome differences. Therefore, we may be missing some signatures of shared functional genes/pathways in our comparative RNA-seq analyses. However, we expect that, using our approach, we detected the most strongly differentially expressed genes in each species, and that those genes play a role in defining and executing the alternate developmental programs within each species. Thus, the comparison between species enables us to begin describing the shared and unique elements of the developmental genetics underlying sex-limited polymorphism. Future work leveraging single-cell approaches is poised to allow a finer-scale dissection of the molecular parallels between these polymorphic species.

## METHODS

### Butterfly care and development staging

Butterfly pupae were provided by butterfly breeders in the Philippines and grown in the greenhouse at the University of Chicago. After emergence, adult butterflies were separated into male and female cages. We collected a single leg from each live adult and used it to genotype *dsx* alleles with custom TaqMan assay probes (Thermo Scientific). Genotype results were used to set up single or multi-pair homozygous crosses in mesh cages with *Citrus* shrubs. From the cages, we collected prepupae each morning and stored them in an incubator matching their natural environment: 70% relative humidity, 25°C, and 16hr light : 8hr dark cycle. P0 pupae were defined as being between 12-24 hours after pupation.

### RNAi

We knocked down *dsx* expression using the RNAi protocol from Ando and Fujiwara 2013 (58) with modifications described in VanKuren et al. 2023 (28). Prepupae were collected from the cross cages in the greenhouse, and pupation was closely monitored so that RNAi experiments could be carried out at the time of pupation when the cuticle was still labile. For injections, we designed dicer substrate siRNAs (DsiRNAs) using IDT’s DsiRNA design tool and a *P. lowii* genome (see below), identifying off-target sites with primer-BLAST (Supplementary Table S1). We injected 2uL of 100uM DsiRNA or 1X PBS as a control into the left hindwing between vein landmarks CU1 and M3. To improve cell permeability for DsiRNA uptake, we covered the injection site with PBS and electroporated the area. The hindwing cuticle cover was then folded back, and the injected pupae were placed into a petri dish with moist paper towels. We stored the petri dishes in the same incubator with settings described above and pinned and imaged the adult butterflies.

### Antibody staining

We characterized DSX spatial expression pattern in *P. lowii* using a polyclonal antibody raised against the closely related species *P. alphenor* (28). For staining, hindwings across different developmental stages were dissected out in 1x RT PBS following the protocol described in VanKuren et al 2023 (28). We used the rabbit anti-Dsx antibody at a 1:250 dilution and co-stained it with either DAPI or Hoescht as a control. To image the dorsal surface of whole mounted wings, a 20x objective and a Z-stack/tile scan was used on a Zeiss LSM 710 confocal microscope at the University of Chicago. Images were then converted to maximum intensity projection and stitched in Zen. We initially processed and scaled images in Fiji and then imported to Inkscape for brightness and contrast adjustment.

### RNA-seq sampling, extraction, and sequencing

We collected three replicates per developmental stage, sex, and *dsx* genotype combination for RNA-seq. Pupae were collected from the incubator at approximately the same time each day to reduce developmental variance between biological replicates. Each replicate consisted of two hindwings dissected from a single pupa and stored in RNAlater (Ambion) in -80°C until extraction. We extracted total RNA using TRIzol (Ambion). TruSeq library preparation using poly-A selection and sequencing (PE100) on an NovaSeq X was done by the Functional Genomics Facility at the University of Chicago (Supplementary Table S2).

### *Papilio lowii* genome sequencing, assembly, and annotation

We extracted HMW gDNA from the thorax of a freshly killed *P. lowii* female homozygous for the mimetic *dsx* allele using the QIAgen GenomicTip G-100 kit. Extractions followed the manufacturer’s instructions, except we incubated chopped fresh tissue in lysis buffer overnight in a thermomixer at 50°C and 200 rpm before purification. We then constructed Oxford Nanopore sequencing libraries using the ONT Ligation Sequencing Kit (LSK-110) and eliminated fragments <10 kb using the PacBio SRE XS kit before sequencing on a MinION Mk1b and R9.4.3 flow cells to 30X - 40X coverage.

We called bases using Guppy and super high-quality base calling (dna_r9.4.1_450bps_sup.cfg), then assembled the genome using these raw reads and Flye v2.9.1 with default settings with expected genome size set to 250 Mb. The initial Flye assemblies were each polished using the Guppy basecalls and Medaka v1.7.2 (medaka_consensus) with the appropriate error model (r941_min_sup_g507). We then purged duplicates using purge_dups v1.2.5 (59). We used a custom repeat library for *Papilio alphenor* (28), the RepBase 20181026 “arthropoda” database, and Dfam 20181026 database to identify and mask repeats genome-wide in both assemblies using RepeatMasker. The final assembly comprised 251.9 Mb in 751 scaffolds (N50 1.6 Mb). BUSCO v5 using the OrthoDBv10 endopterygota database showed the assembly contained 98.7% complete (0.5% duplicated), 0.1% fragmented, and 1.2% missing SCOs.

We annotated the *P. lowii* assembly using EvidenceModeler 1.1.1 (60). We first assembled a high-quality transcript database using PASA (60) and the PE100 data generated here. After adapter trimming, we performed *de novo* using Trinity v2.10.0 (61) and genome-guided assembly using StringTie v1.3.3 (62). RNA-seq data were also mapped to the assembly using STAR 2.6.1d (63), and the resulting alignments used to generate genome-guided assemblies with Trinity and StringTie v1.3.3 (62). We combined *de novo* and genome-guided assemblies using PASA 2.4.1 (64). Evidence for protein-coding regions came from mapping the UniProt/Swiss-Prot (2020_06) database and all Papilionoidea proteins available in NCBI’s GenBank nr protein database (downloaded 6/2020) using exonerate (65). We identified high-quality multi-exon protein-coding PASA transcripts using TransDecoder (transdecoder.github.io), then used these models to train and run Genemark-ET 4 (66) and GlimmerHMM 3.0.4 (67). We also predicted gene models using Augustus 3.3.2 (68), the supplied *heliconius_melpomene1* parameter set, and hints derived from RNA-seq and protein mapping above. Augustus predictions with >90% of their length covered by hints were considered high-quality models. Transcript, protein, and *ab initio* data were integrated using EVM with the weights in Supplementary Table S3.

Raw EVM models were then updated twice using PASA to add UTRs and identify alternative transcripts. Gene models derived from transposable element proteins were identified using BLASTp and removed from the annotation set. We manually curated the *dsx* region. The final annotation comprises 21,733 genes encoding 30,844 protein-coding transcripts, containing 97.6% complete and missing 1.5% of endopterygota single-copy orthologs according to BUSCO v5 and OrthoDB v10. We functionally annotated protein models using eggNOG’s emapper-2.0.1b utility and the v2.0 eggNOG database (69).

### Differential expression analysis

We used the annotated genome to quantify transcript expression levels in Salmon v1.10.0 with bias correction (70). Quantification results were imported and normalized in R using tximport v1.34, and filtered for lowly expressed genes, which were defined as genes with an average normalized count below 5 across all samples. From filtered, normalized data, we used mapping rate, classical PCA (95% CI), and robust PCA using the PcaGrid function in rrcov v1.7-6 (71) to identify outlier samples. After removing outliers, we used filtered, normalized gene-level quantification data in DESeq2 v1.46 to identify differentially expressed genes at each stage in pairwise comparisons of the female groups (mimetic female versus non-mimetic females). We also used maSigPro v1.78 (72) to identify differentially expressed genes, defined as genes with significantly different temporal expression profiles in mimetic females compared to all other genotype-sex groups (non-mimetic females, mimetic males, non-mimetic males). Significant genes were first selected using a q-value cutoff of 0.01 in the p.vector() function. Subsequently, variable selection was performed based on a p-value threshold of 0.05 in T.fit(), and genes with good model fits were defined as those with an R² > 0.8. To identify orthologous genes between the two species, we used BLASTp v2.14.40 and kept the one-to-one reciprocal best hits.

### Co-expression network reconstruction and module preservation

Using RNA-seq data from *P. lowii* generated in this study, along with previously published RNA-seq data from *P. alphenor* (28), we constructed hindwing developmental gene co-expression networks for both species. For each species separately, networks were built using WGCNA v1.73 (45) with normalized gene-level expression data, and both adjacency and topological overlap matrices were generated using signed Pearson correlation coefficients. To examine the relationships between gene modules and traits (stage, sex, genotype, sex-genotype combination), Pearson correlations were calculated between each module eigengene and trait variable, with corresponding p-values determined using Student’s correlation test. We then tested each module for enrichment or depletion of DEGs using Fisher’s Exact Test; modules with adjusted p-values < 0.01 were considered significantly enriched or depleted in DEGs. We then conducted GO enrichment analysis for each module using topGO v2.58 (73). GO term assignments were derived from eggNOG-mapper v2.1.12 annotations (69). We tested for enrichment of biological process terms using Fisher’s Exact Test, with a minimum node size of 5 genes and an adjusted p-value threshold of 0.01. The top 50 enriched terms were retained for each module. After completing network reconstruction for each species, we used the ModulePreservation function in WGCNA to identify orthologous modules between both species. For our input datasets to test for network preservation across species, we used the full normalized expression data and the gene assignments to modules from each species.

## Acknowledgements

We thank the University of Chicago greenhouse staff and the University of Chicago Functional Genomics Facility (RRID:SCR_019196) for research support, the University of Chicago’s Center for Research Informatics for computational support, and Nipam Patel for providing the 4F11 Engrailed antibody. We also appreciate lab members Darli Massardo, Wei Lu, Paula Fernandez-Begne, Hsiang-Yu Tsai, and William Zhang for valuable feedback and discussion on the manuscript. This work was supported by the GME NIGMS T32 training grant and an ARCS Foundation award to SIS, the Chicago Fellows program to MMD, and NIH R35 GM131828 to MRK.

## Author contributions

SIS - Conceptualization, Investigation, Visualization, Writing - original draft, review & editing; MMD - Conceptualization, Investigation; NWV - Conceptualization, Investigation, Writing - review & editing; PH - Investigation; MRK - Conceptualization, Funding acquisition, Supervision, Writing - review & editing

## Data Availability

Illumina RNA sequencing data and the genome assembly and annotation are publicly available in the National Center for Biotechnology Information (NCBI) under BioProject PRJNA1230174. Anti-DSX antibody is available from the authors upon request.

