## Supplementary material for "A shared gene but distinct dynamics regulate mimicry polymorphisms in closely related species": figS1.pdf

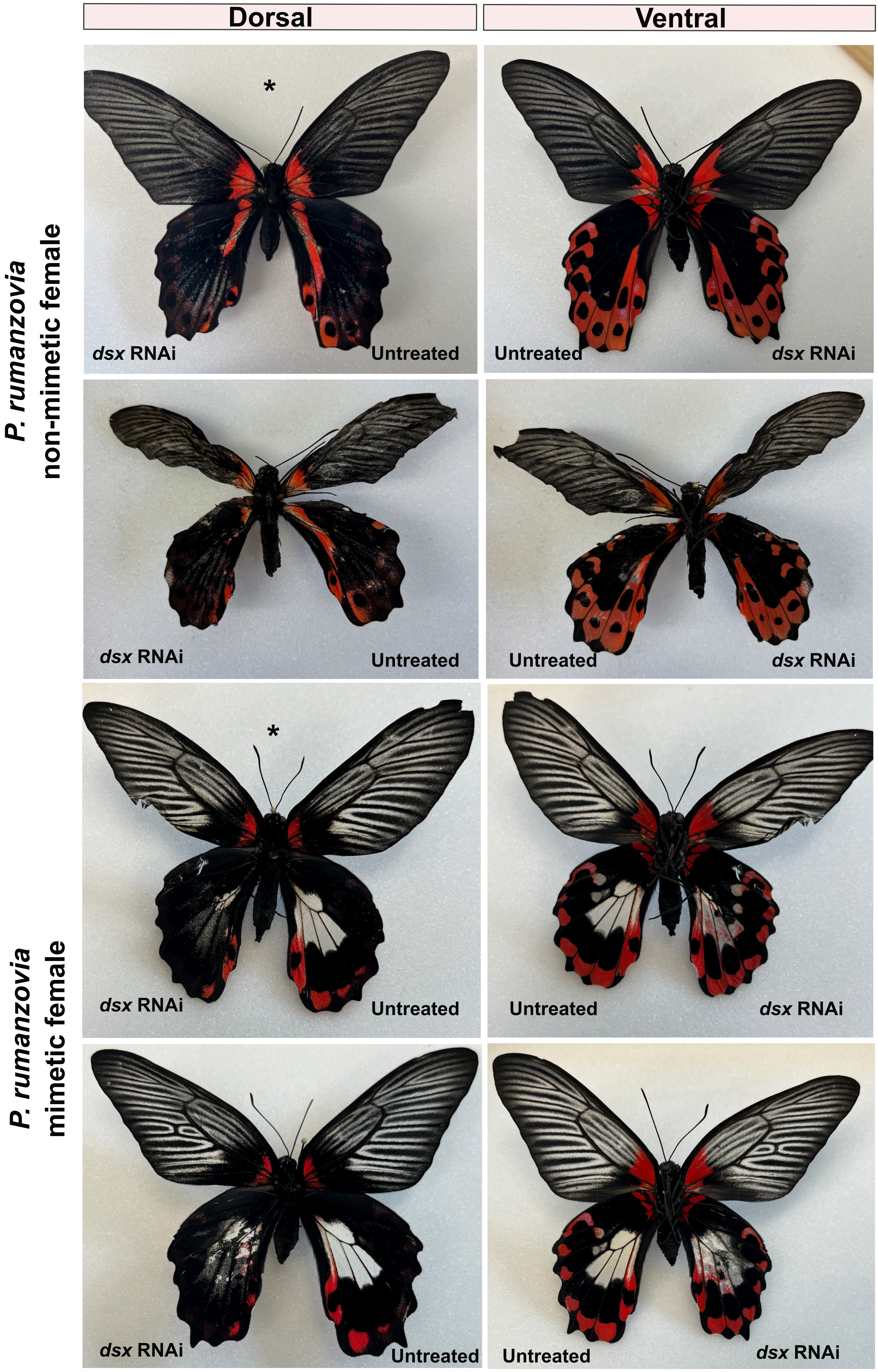
Dorsal
Ventral
P. rumanzovia
non-mimetic female
P. rumanzovia
mimetic female
*
dsx RNAi
Untreated
Untreated
dsx RNAi
dsx RNAi
Untreated
Untreated
dsx RNAi
*
dsx RNAi
Untreated
Untreated
dsx RNAi
dsx RNAi
Untreated
Untreated
dsx RNAi

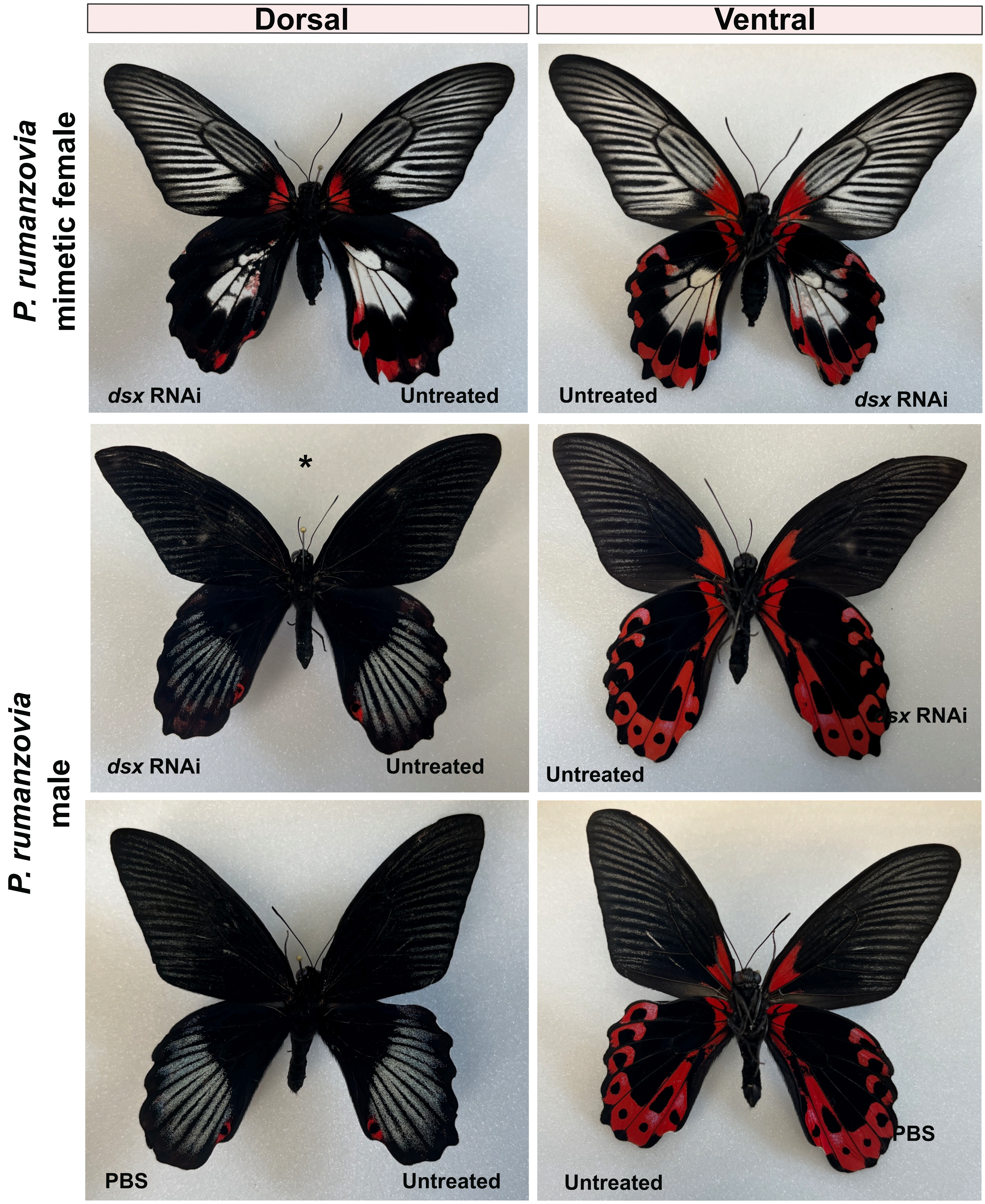

Dorsal
Ventral
P. rumanzovia
mimetic female
dsx RNAi
Untreated
Untreated
dsx RNAi
P. rumanzovia
male
*
dsx RNAi
Untreated
sx RNAi
Untreated
PBS
Untreated
PBS
Untreated

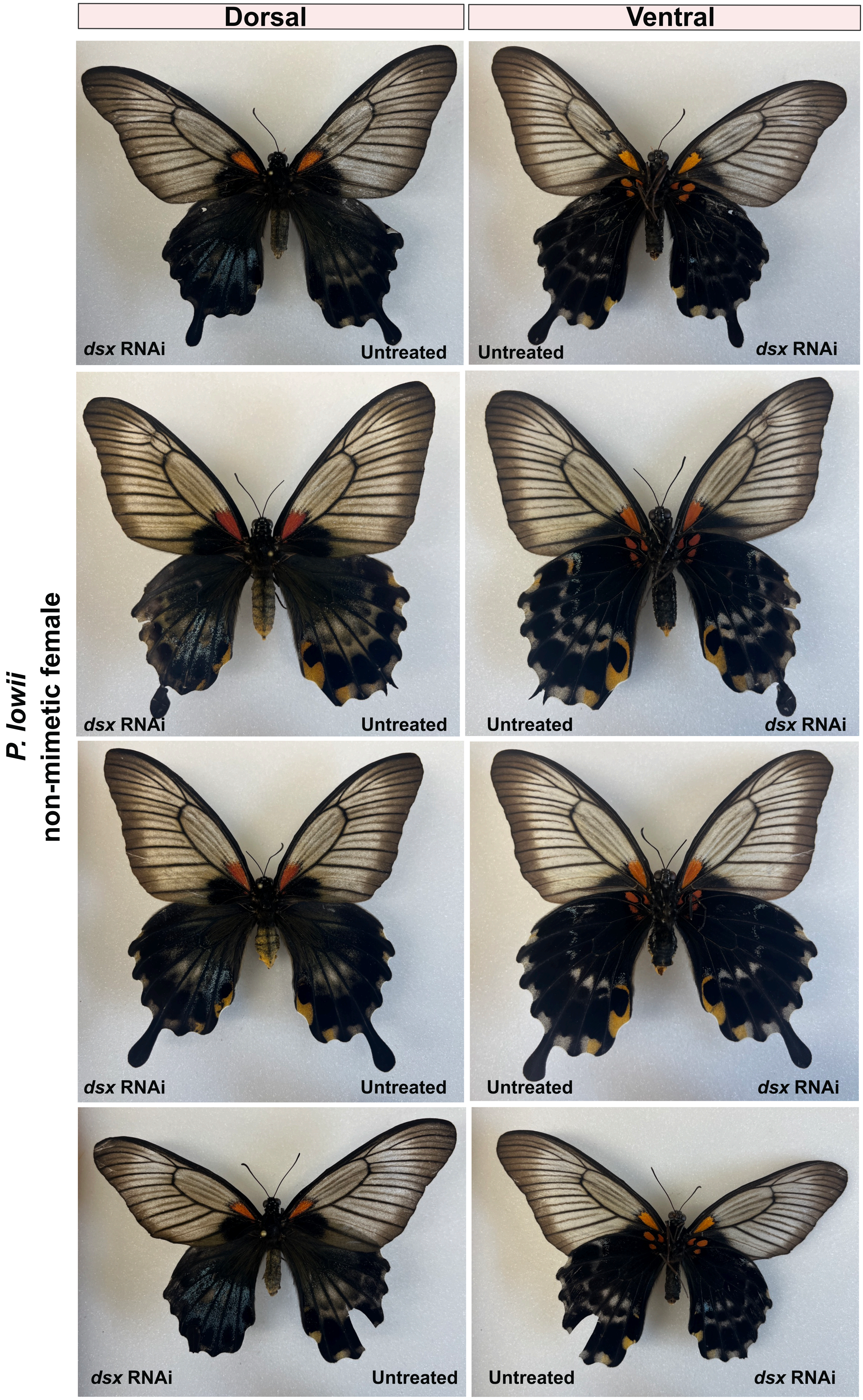

Dorsal
Ventral
P. lowii
non-mimetic female
dsx RNAi
Untreated
Untreated
dsx RNAi
dsx RNAi
Untreated
Untreated
dsx RNAi
dsx RNAi
Untreated
Untreated
dsx RNAi
dsx RNAi
Untreated
Untreated
dsx RNAi

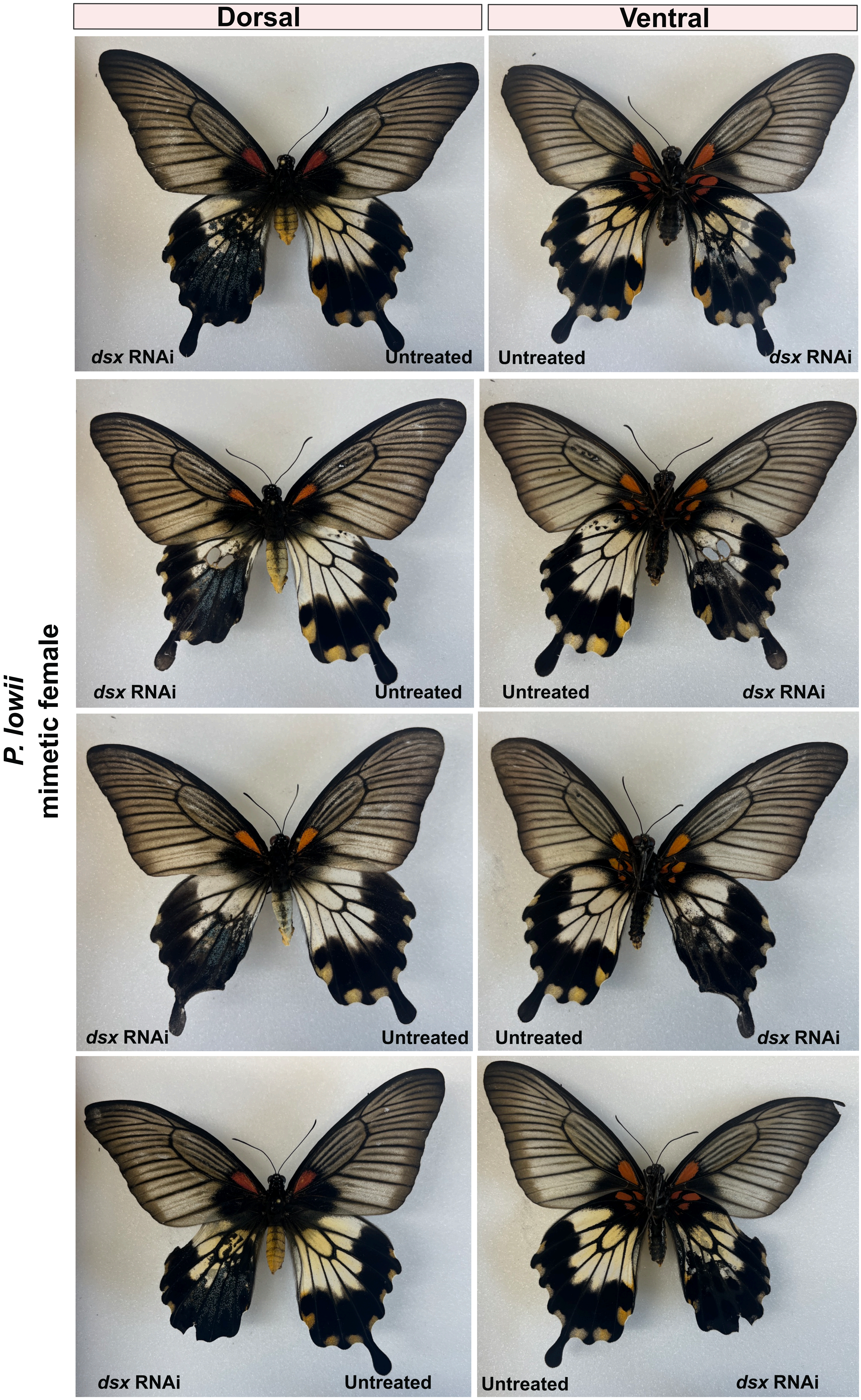

Dorsal
Ventral
P. lowii
mimetic female
dsx RNAi
Untreated
Untreated
dsx RNAi
dsx RNAi
Untreated
Untreated
dsx RNAi
dsx RNAi
Untreated
Untreated
dsx RNAi
dsx RNAi
Untreated
Untreated
dsx RNAi

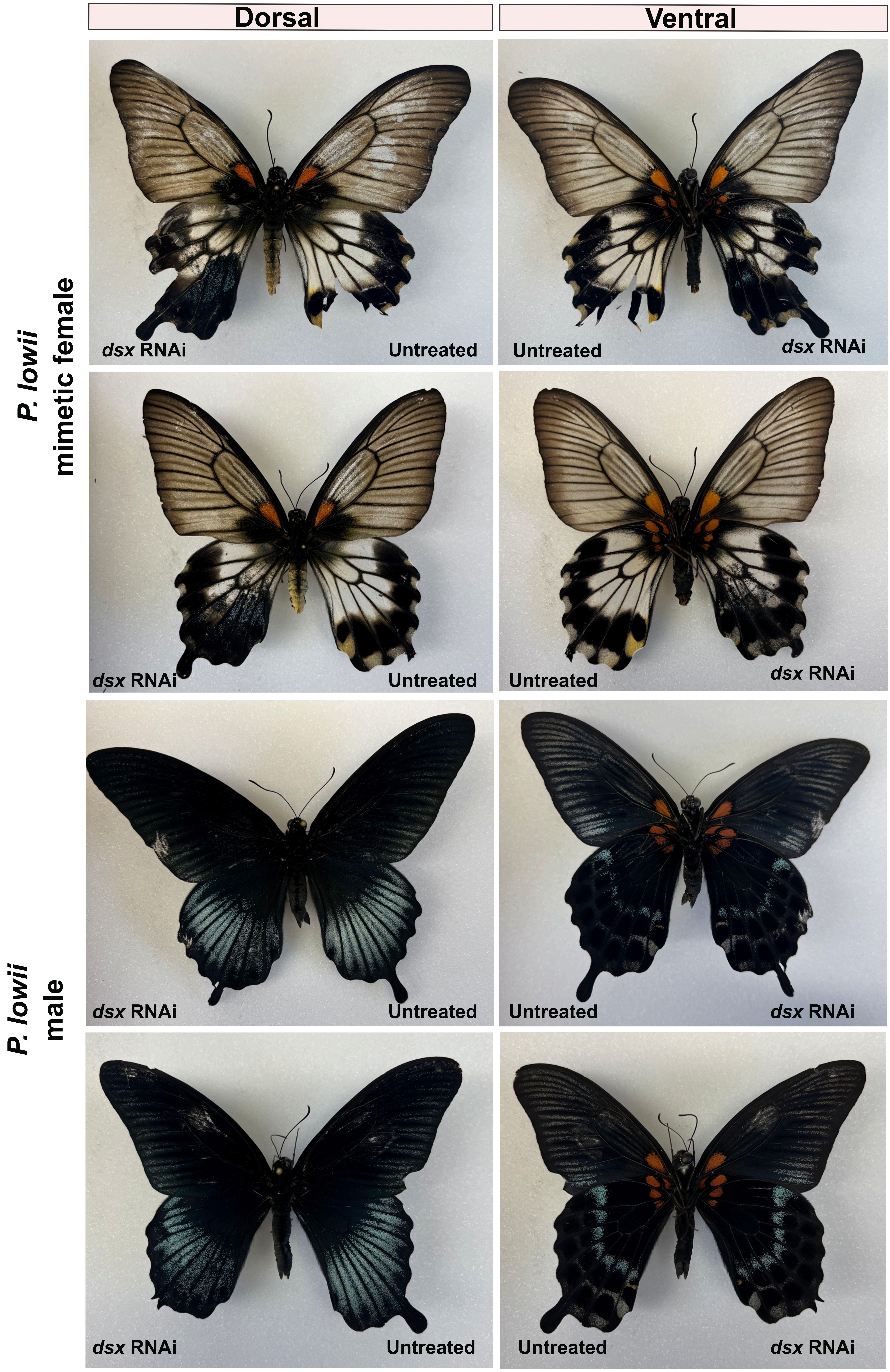

Dorsal
Ventral
P. lowii
mimetic female
P. lowii
male
dsx RNAi
Untreated
Untreated
dsx RNAi
dsx RNAi
Untreated
Untreated
dsx RNAi
dsx RNAi
Untreated
Untreated
dsx RNAi
dsx RNAi
Untreated
Untreated
dsx RNAi

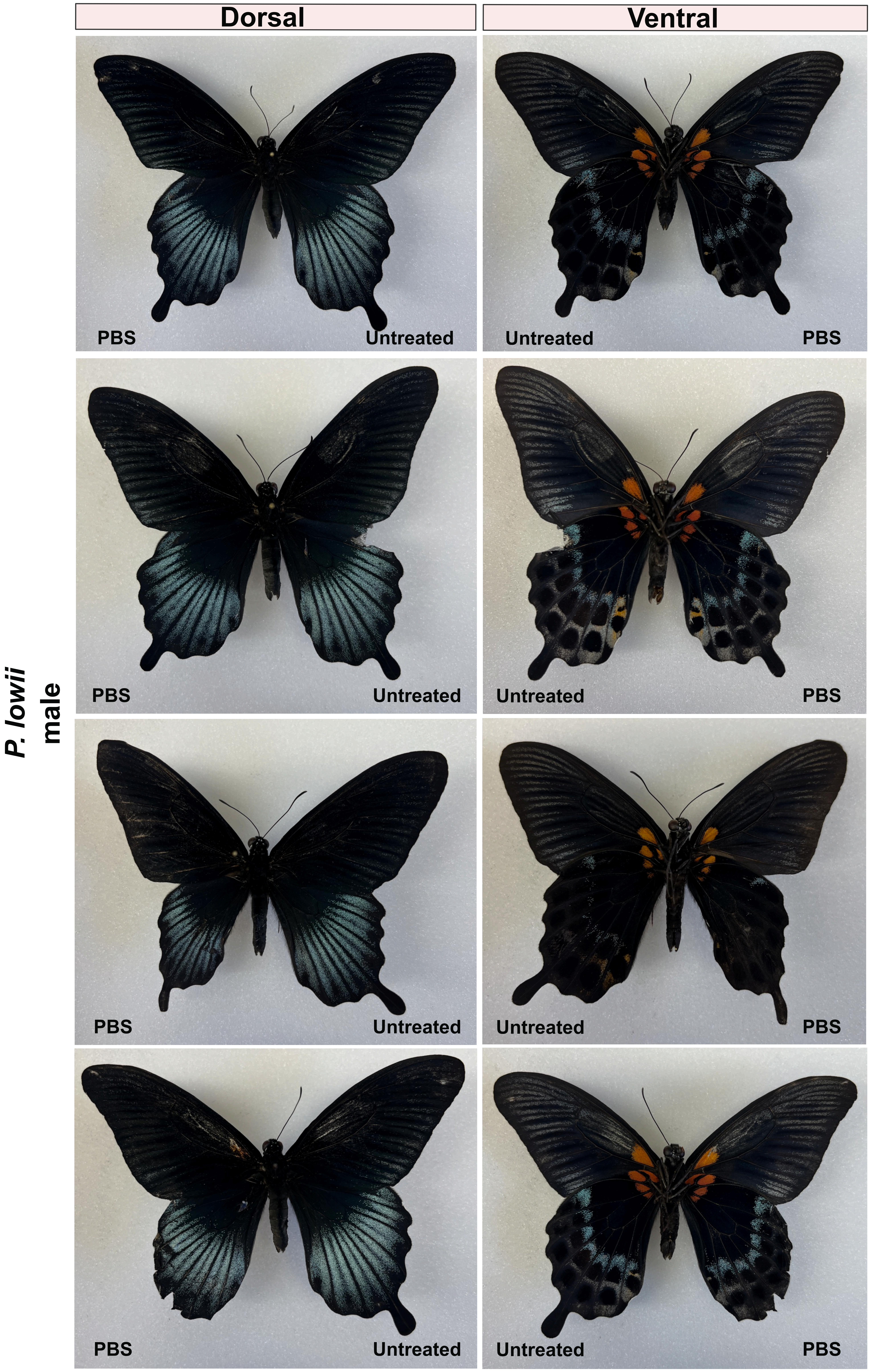
Dorsal
Ventral
P. lowii
male
PBS
Untreated
Untreated
PBS
PBS
Untreated
Untreated
PBS
PBS
Untreated
Untreated
PBS
PBS
Untreated
Untreated
PBS

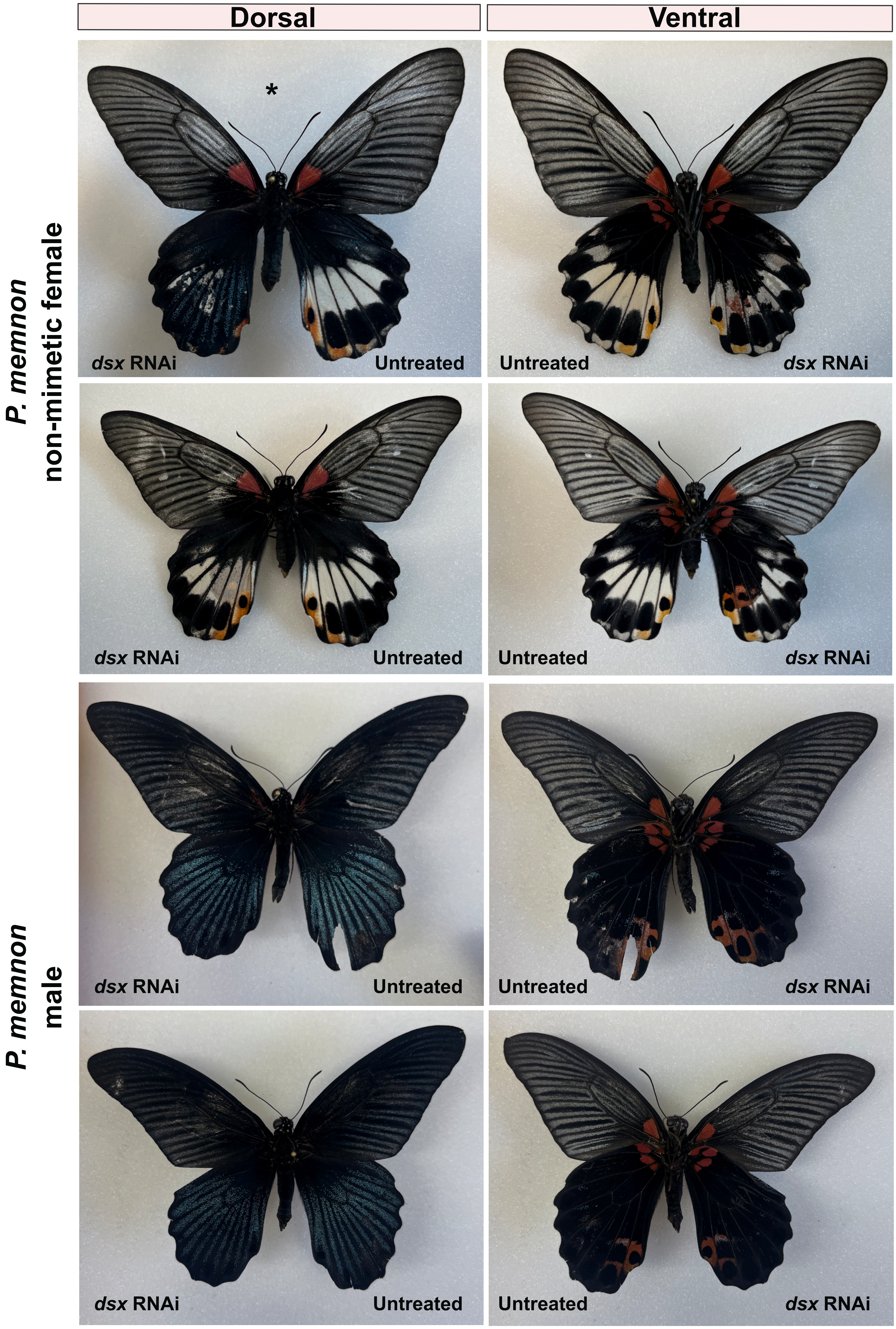

Dorsal
Ventral
P. memnon
non-mimetic female
P. memnon
male
*
dsx RNAi
Untreated
Untreated
dsx RNAi
dsx RNAi
Untreated
Untreated
dsx RNAi
dsx RNAi
Untreated
Untreated
dsx RNAi
dsx RNAi
Untreated
Untreated
dsx RNAi

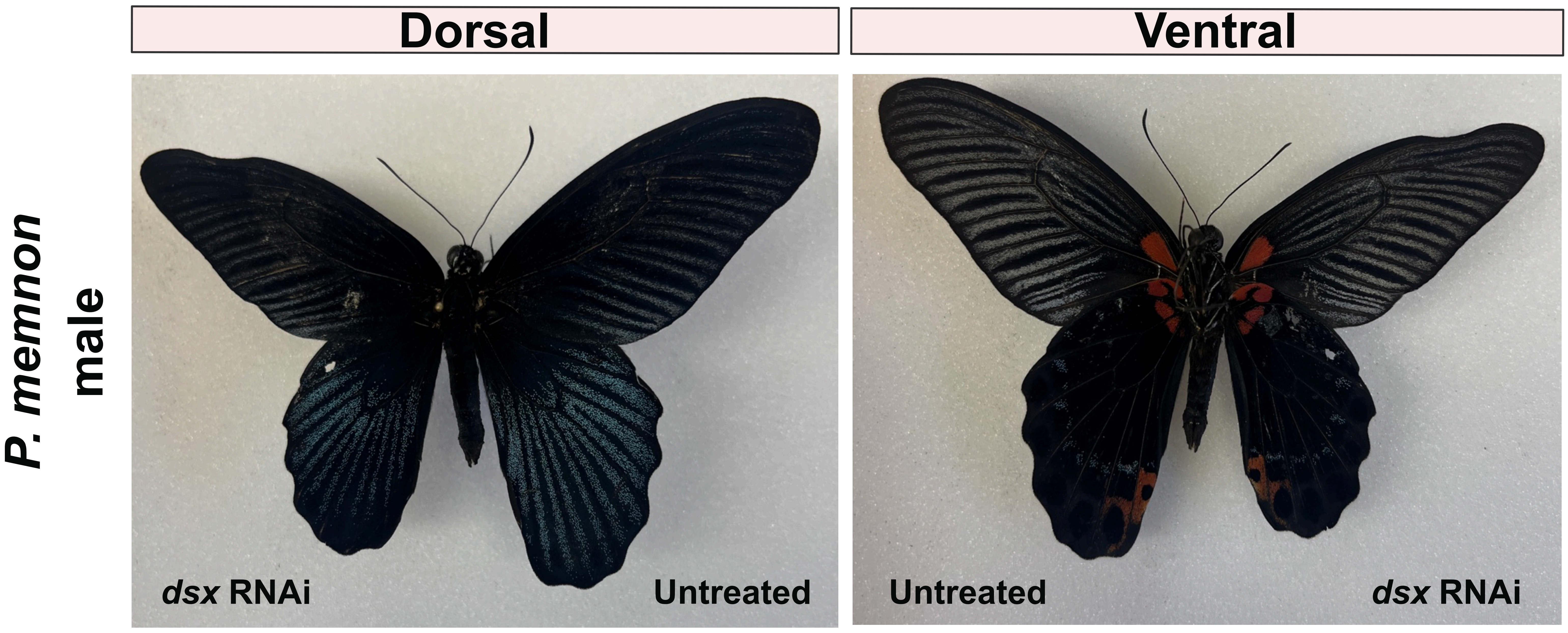


**Figure S1. Dorsal and ventral image of additional RNAi treated individuals.** Pupae were injected with 1.5 uL 100 uM DsiRNAs or PBS as a control. Individuals with * on the dorsal are shown in Figure 1, and included here to show the ventral surface.
