## Supplementary material for "A shared gene but distinct dynamics regulate mimicry polymorphisms in closely related species": supp_figs_S2-S9.pdf

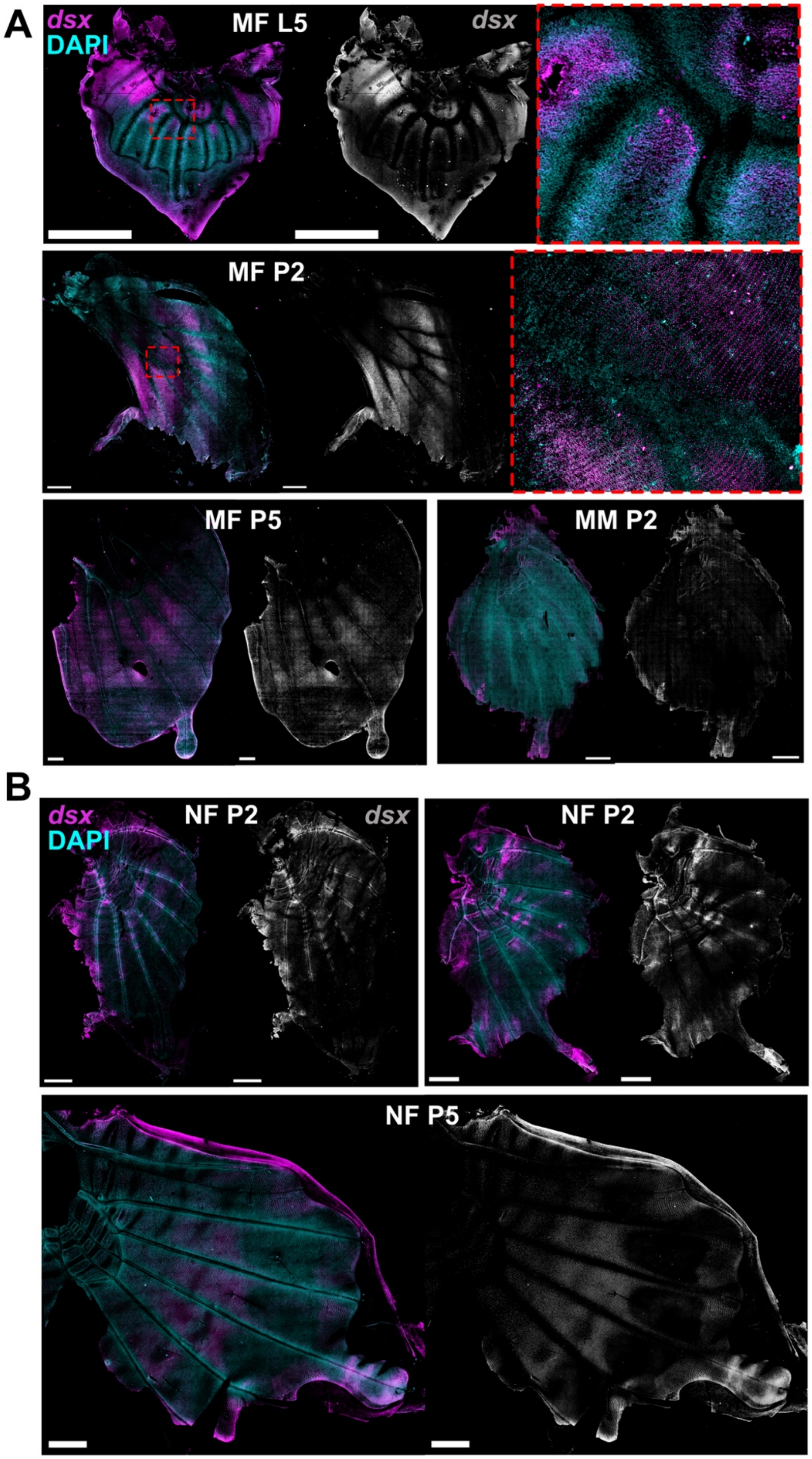


**Figure S2. Additional anti-Dsx stains in pupal *P. lowii* hindwings. A)** Mimetic wings; insets show enrichment near discal cell and localization to scale cells by 2 days post-pupation. **B)** non-mimetic females; Dsx prefigures the light patch by 5 days post-pupation.

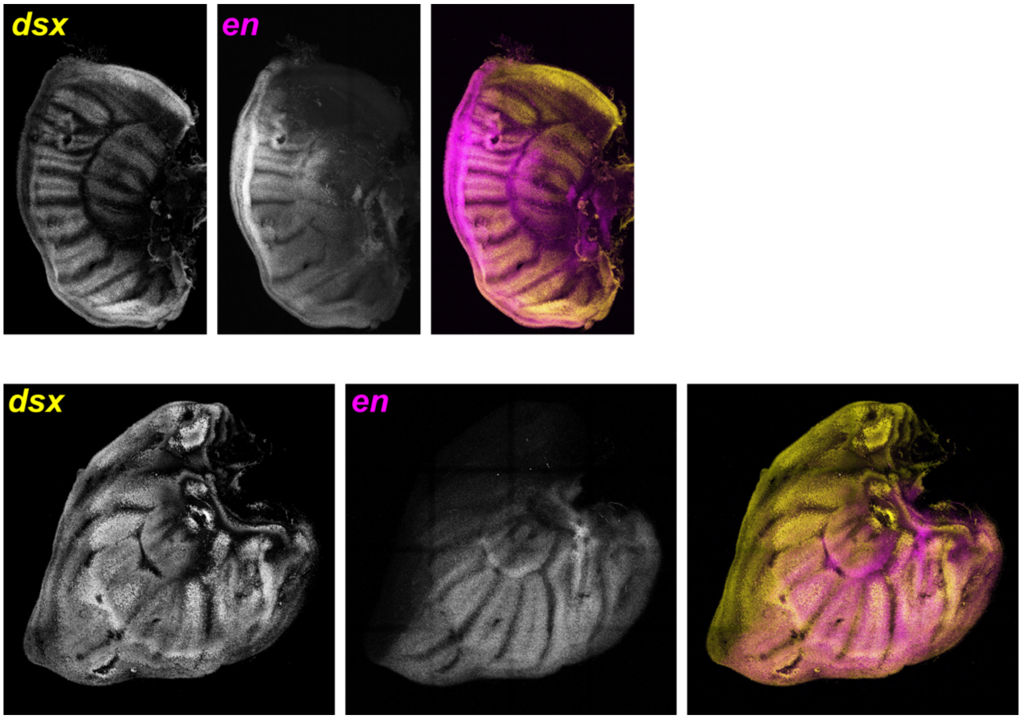


**Figure S3. Anti-dsx stain in late larval *P. alphenor* wings**. Forewing images on top and hindwing on bottom, using hybridization chain reaction following HCR In Situ Protocol V.1 by Bruce et al. 2021 (dx.doi.org/10.17504/protocols.io.bunznvf6) with *doublsex* and *engrailed* probes designed by Molecular Instruments. *dsx* shows uniform expression across the hindwing.

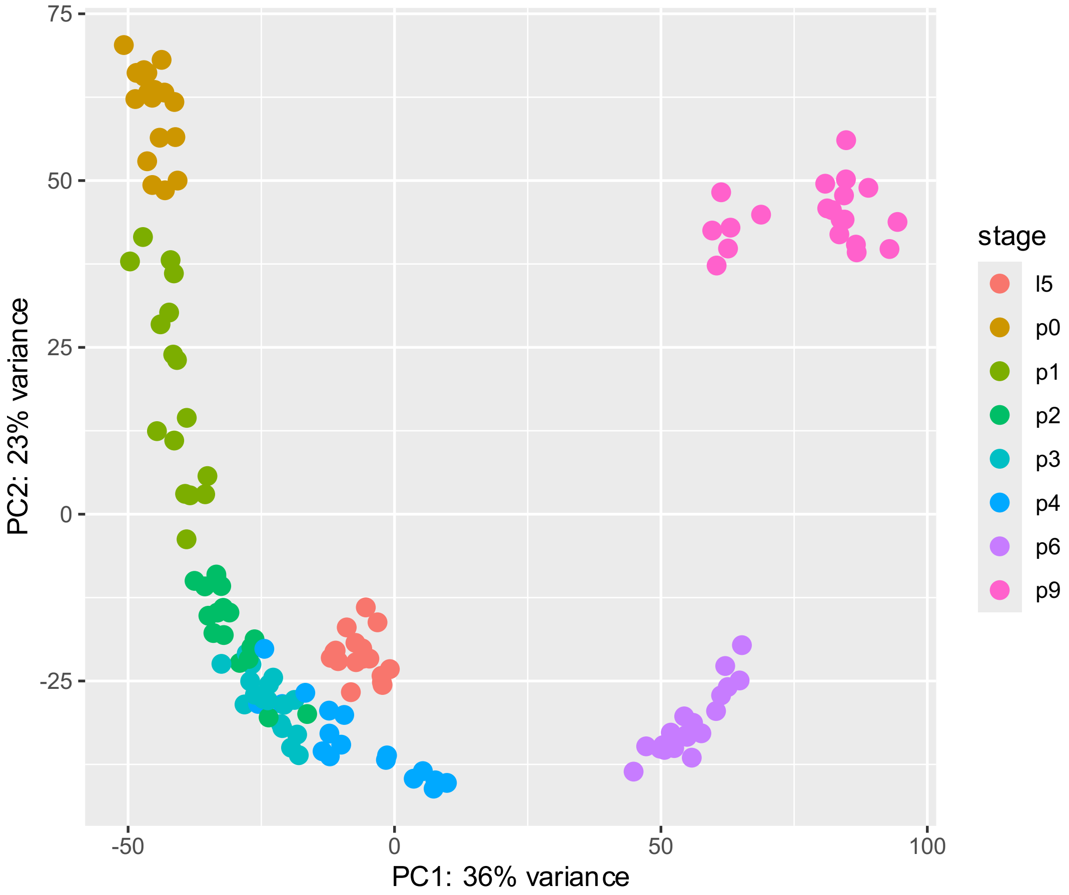


**Figure S4. RNA-seq sample clustering.** PCA of filtered RNA-seq samples based on normalized expression data.

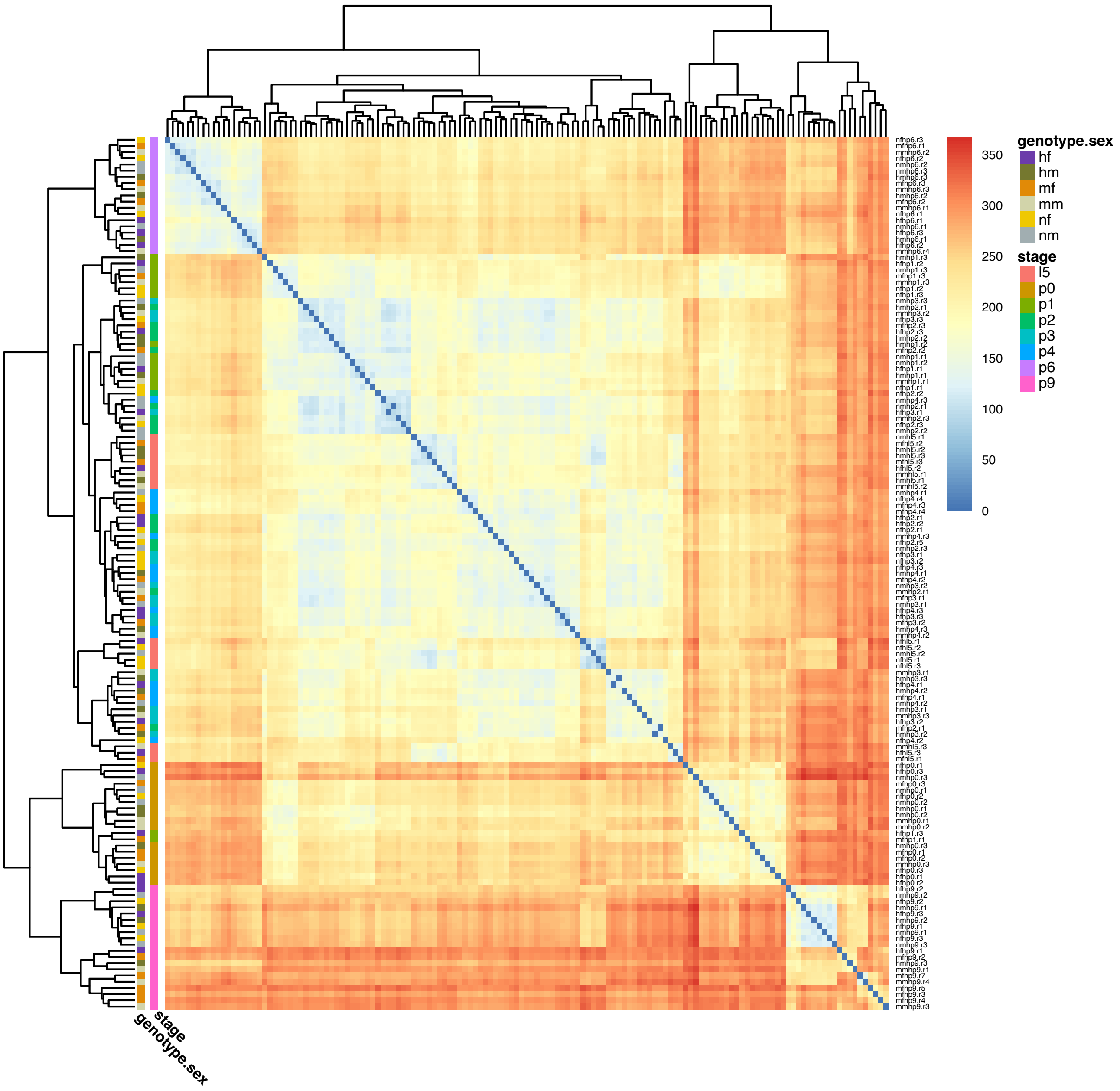


**Figure S5. RNA-seq sample clustering.** Heatmap of filtered RNA-seq samples based on normalized expression data, clustered using the pheatmap package and *k* means clustering.

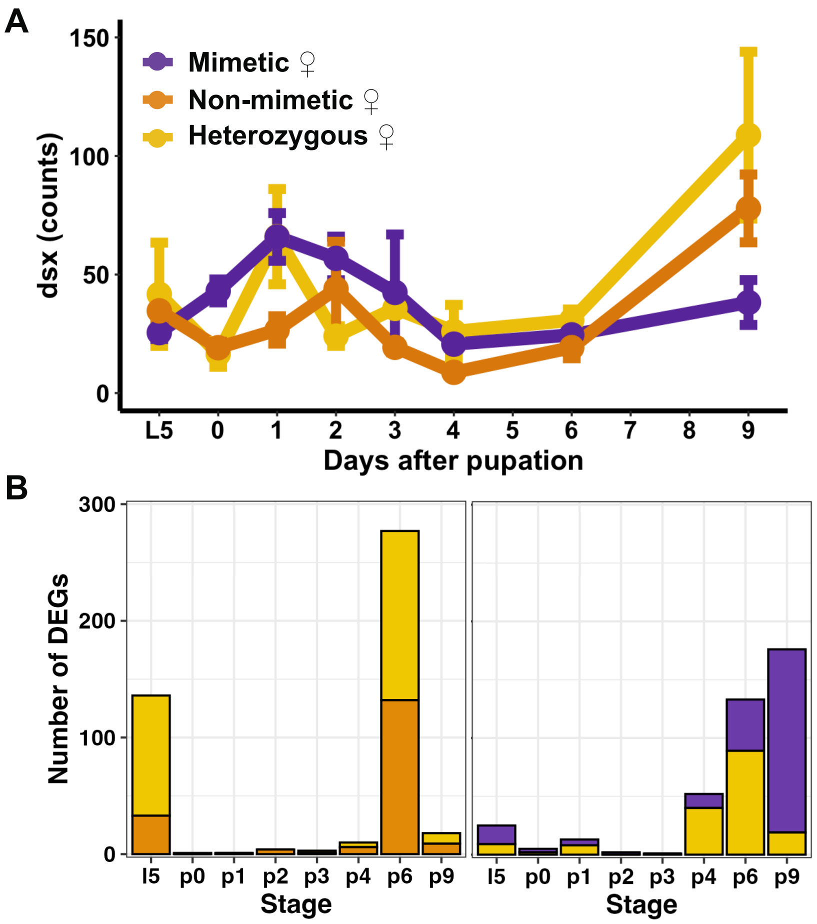


**Figure S6. RNA-seq results in heterozygous females. A)** *doublesex* expression in all female groups. **B)** Genes differentially expressed and upregulated in heterozygous females compared to non-mimetic females (left) and heterozygous females compared to mimetic females (right).

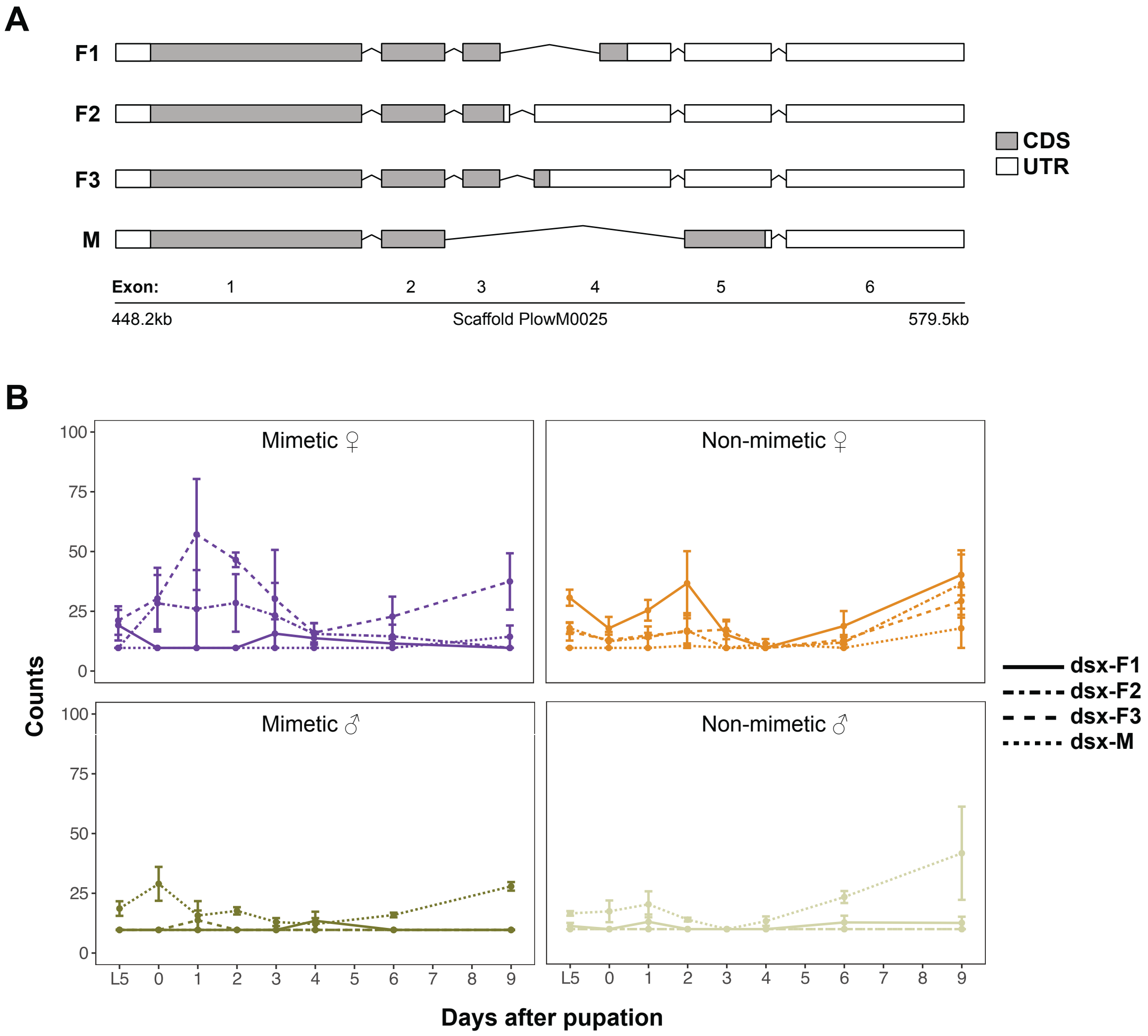


**Figure S7. *Doublesex* isoform expression. A)** Exon-intron structures of four *dsx* isoforms in *P. lowii* based on the annotated genome. Introns not to scale. **B)** Normalized transcription expression levels in each sex-genotype group

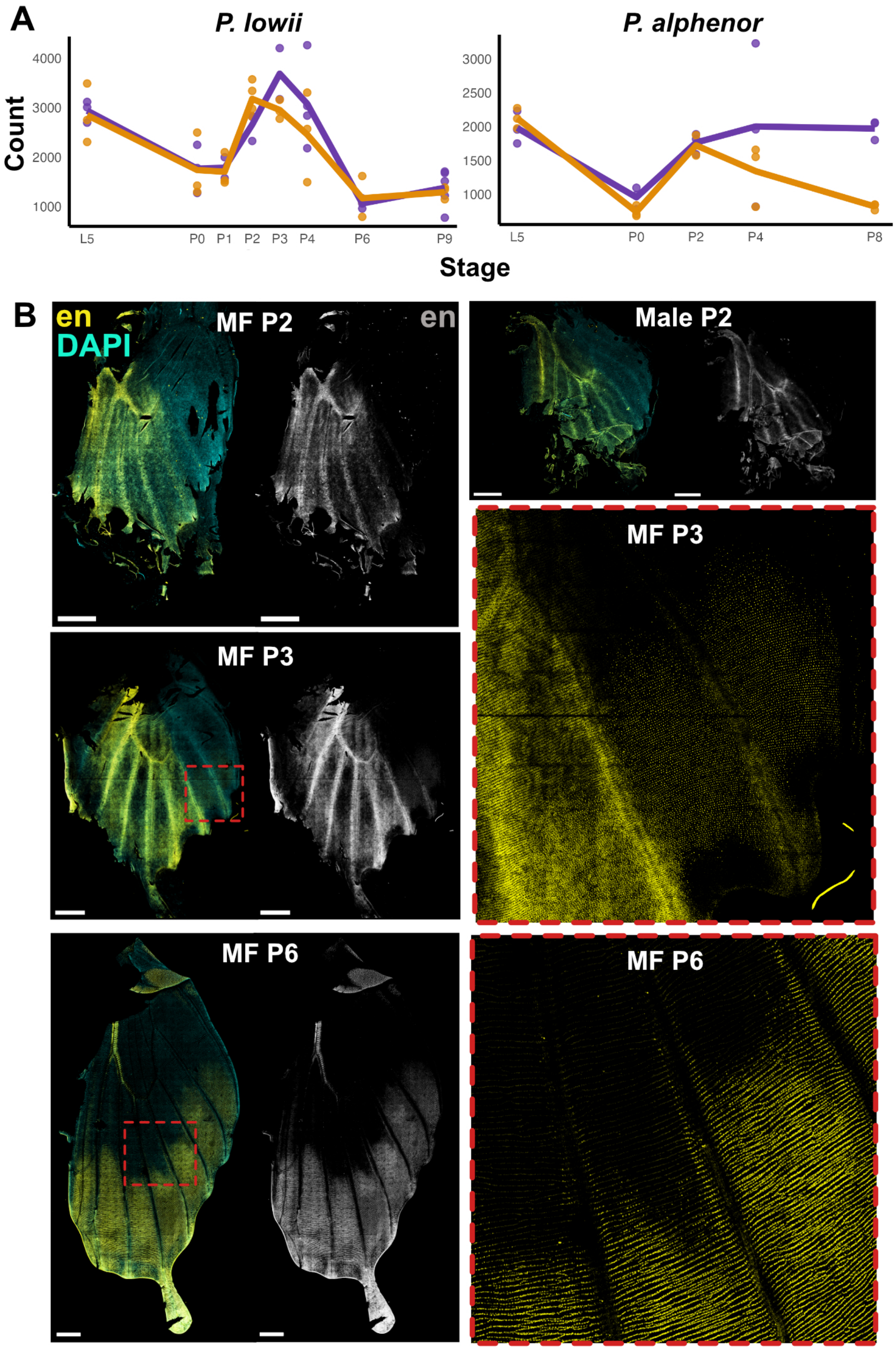


**Figure S6. *Engrailed* expression in pupal hindwings. A)** Counts of *en* expression over the developmental time series in *P. lowi* (not DE) and *P. alphenor* (DE). **B)** Anti-En staining in mimetic female hindwings from early to mid-pupal development. During early pupal development, En performs its ancestral function of AP patterning, however by 3 days post-pupation it is expressed in an additional domain, likely beginning to specify the melanic regions of the adult wing as observed in *P. alphenor* (VanKuren et al 2023). By 6 days post pupation, En expression is restricted to the melanic regions of the mimetic wing color pattern.

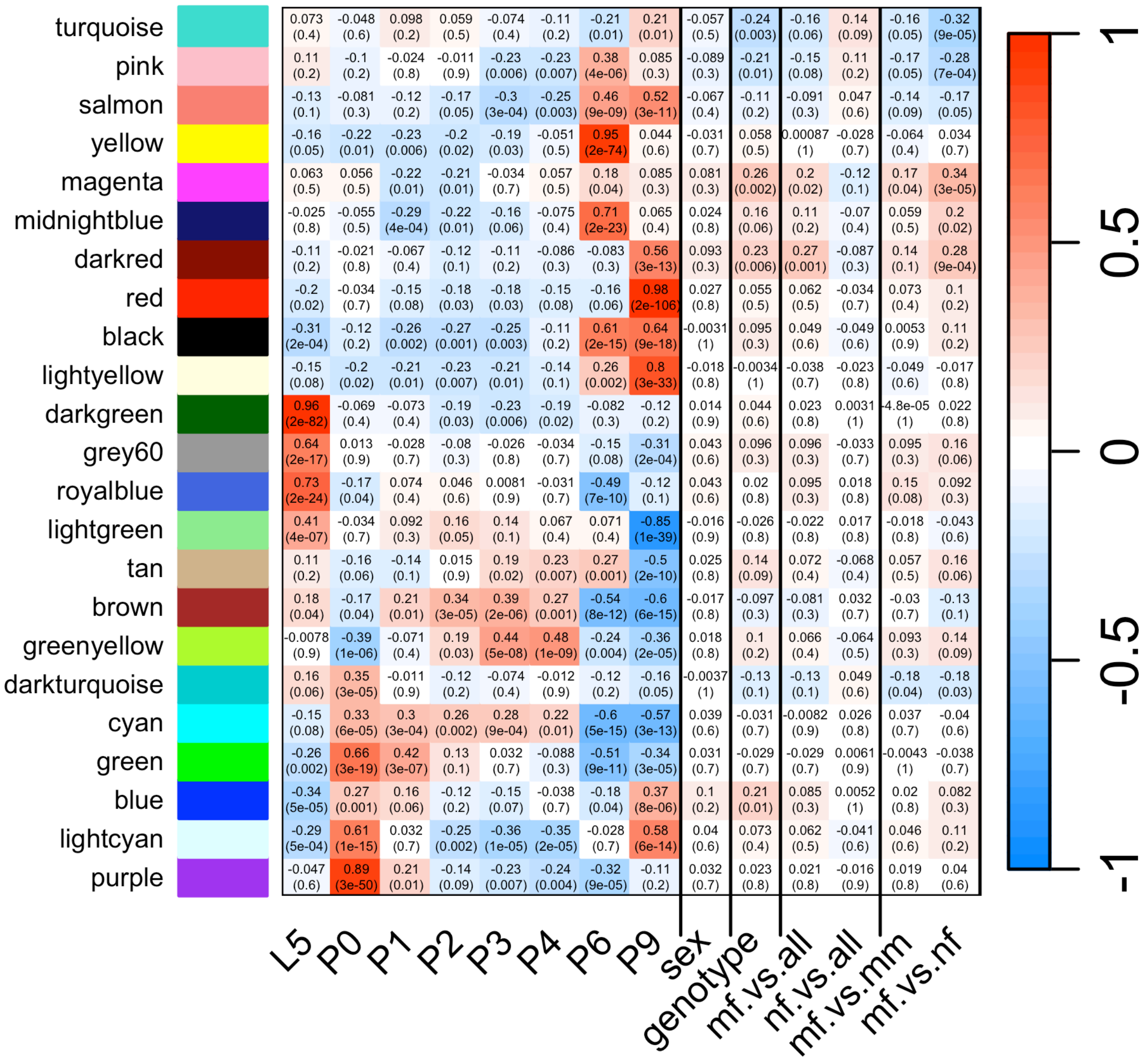


**Figure S9. Module-trait relationships in *P. lowii***. In each grid, the top number is the correlation between module eigengenes and traits, and the bottom number in parentheses is the bonferroni-corrected p-value.

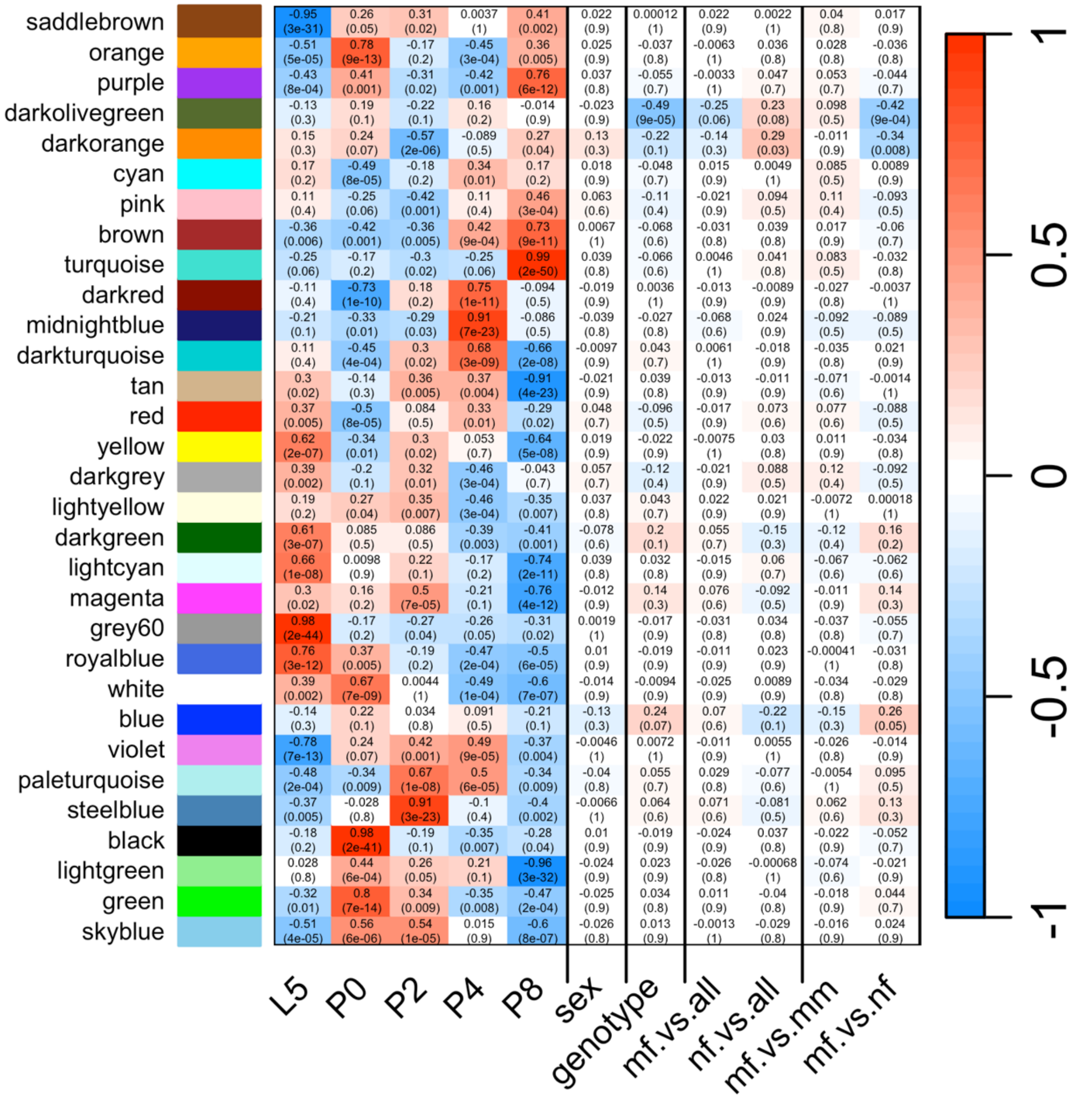


**Figure S10**. **Module-trait relationships in *P. alphenor.*** In each grid, the top number is the correlation between module eigengenes and traits, and the bottom number is the bonferroni-corrected p-value.

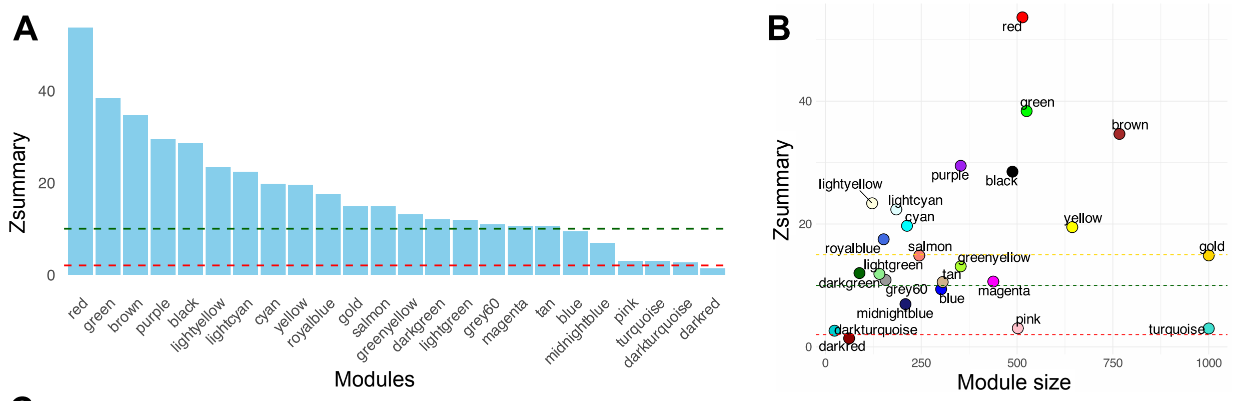


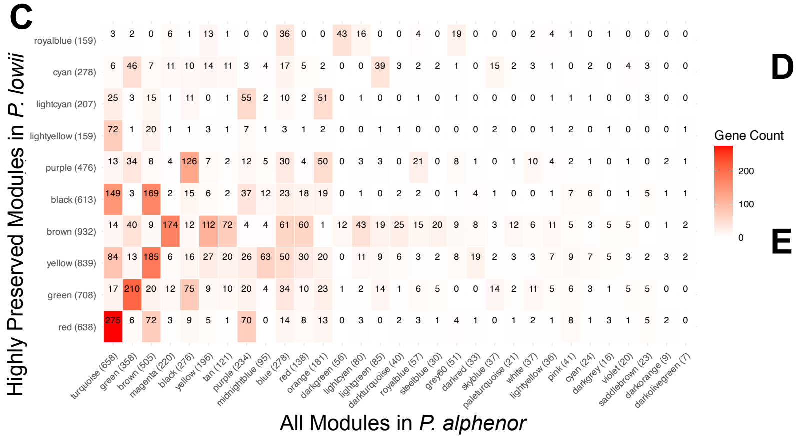


**D**

| | Module in... | | | Enriched for DEGs in... (adjusted p < 0.05) | |
|---|---|---|---|---|---|
| | *P. lowii* | *P. alphenor* | BH adjusted p | *P. lowii* | *P. alphenor* |
| 1 | Red | Turqoise | 7.45E-123 | Yes (E) | No |
| 2 | Green | Green | 1.39E-123 | **Yes (E)** | **Yes (E)** |
| 3 | Brown | Magenta | 9.11E-103 | **Yes (D)** | **Yes (D)** |
| 4 | Purple | Black | 1.32E-75 | Yes (E) | No |
| 5 | Royalblue | Darkgreen | 3.91E-54 | **Yes (D)** | **Yes (D)** |
| 6 | Black | Brown | 3.44E-54 | No | Yes (D) |

**E**

| | |
|---|---|
| 1 | Fatty acid & metabolic processes, oxidative phosphorylation, nucleotide biosynthesis |
| 2 | Toll signaling, immune response |
| 3 | Cell cycle and division |
| 4 | metal ion response, catabolism, hormone regulation |
| 5 | RNA processing, gene expression regulation |
| 6 | Vesticle transport, protein localization, secretion |

**Figure S11. WGCNA module preservation and orthology. A)** Distribution of Zsummary scores, which are a composite statistic of module preservation across reference species (plow) and test species (palp). Red line indicates Zscore of no preservation (< 2), green line indicates weak or moderate preservation (< 9), and Zscores about the green line indicate strong preservation. **B)** Correlation between Zsummary statistic and module size. The gold line indicates the Z-score of the control module (gold); this module is made by randomly sampling 1000 genes and testing the preservation of this module. A high Zsummary indicates strong network preservation generally, and we focused on modules with Zsummary value greater than the gold. **C)** Gene membership overlap between highly preserved plowii modules and all modules in alphenor to identify orthologous ones. The module sizes are indicated in parenthesis, however they may be biased in *P. alphenor* due to the number of orthologous gene IDs we could assign between both species. **D)** Most likely orthologous modules between species and whether or not those modules are enriched for DEGs (D = deficient in DEGs, and E = enriched, that is, greater or fewer DEGs observed in a module than expected by chance). **E)** Summary of top GO terms and processes in the likely orthologous modules in D.
